# Glutamine-Dependent Slc25a39–Nrf2 Axis Couples Mitochondrial Dynamics with Metabolic Reprogramming to Establish Myogenic Commitment

**DOI:** 10.1101/2025.10.02.680066

**Authors:** Brian Kelly, Chia-Hua Wu, Kaan I. Eskut, Elizabeth McCauley, Sajina Dhungel, Samantha Pavlecic, Grace Bieghler, Jelena Perovanovic, Dean Tantin, Jakub Famulski, Jeramiah Smith, Chintan Kikani

## Abstract

Myogenic commitment is a decisive and irreversible step in skeletal muscle regeneration, necessitating proliferating myoblasts to integrate metabolic cues with nuclear transcriptional programs. Among amino acids, glutamine is uniquely positioned to influence this transition by coupling energy production to macromolecule biosynthesis and epigenetic regulation. We reasoned that myoblasts must sense glutamine availability to ensure orderly progression toward commitment, and we tested this by examining the molecular consequences of acute glutamine withdrawal. We find that continued glutamine oxidation is required to sustain glycolysis, maintain mitochondrial fission, and preserve a redox balance that supports progression towards myogenic commitment. In its absence, myoblasts undergo a reductive shift, characterized by mitochondrial elongation, membrane depolarization, and suppression of glycolysis, ultimately leading to growth arrest. Transcriptomic profiling reveals reduced *MyoD* and *MKi67*, accompanied by increased *Sprouty1* levels, defining a reversible non-proliferative state that resembles but is distinct from quiescent and reserve cells. We term this state Poised Metabolic Arrest (PMA), a cellular response to glutamine limitation during myogenic progression. Mechanistically, PMA is driven by Nrf2-dependent increased glutathione (GSH) biosynthesis and upregulation of mitochondrial GSH carrier Slc25a39 when glutamine is limited. Depleting mitochondrial glutathione or silencing Slc25a39 forces exit from PMA. However, this premature exit compromises subsequent differentiation potential, indicating PMA serves to preserve differentiation competence when glutamine is limited. Consistent with this, both loss and overexpression of Slc25a39 impair myoblast differentiation in vitro and disrupt regeneration in vivo. Together, these data suggest that a reciprocal Slc25a39–Nrf2 redox axis functions as a nutrient-dependent checkpoint, coupling glutamine availability to mitochondrial remodeling and metabolic reprogramming, necessary to establish irreversible myogenic commitment.

## Introduction

Tissue regeneration is orchestrated by metabolic processes that intersect with transcriptional responses to guide stem cells through successive stages, leading to differentiation^1,2^. In skeletal muscle, muscle stem cells (MuSCs), also known as satellite cells, depend on extracellular cues to activate transcriptional programs that drive regeneration^3,4^. A central feature of this process is the remodeling of mitochondrial structure, which allows stem cells to align their bioenergetic state with regenerative demands. In quiescent MuSCs, mitochondria exist as elongated, interconnected networks^5^. These hyperfused mitochondria are likely to be protected against mitophagy under conditions of low oxidative phosphorylation, thereby preserving the mitochondrial pool until activation^6^. Upon MuSCs activation, this morphology shifts toward increased mitochondrial fission. During differentiation, healthy mitochondria fuse to form elongated networks that support the assembly of respiratory supercomplexes, thereby enhancing the efficiency of oxidative phosphorylation and meeting the heightened bioenergetic and biosynthetic requirements of myofibers. Collectively, these cyclical transitions between fission and fusion provide the metabolic flexibility necessary to maintain cellular homeostasis and sustain the regenerative capacity of skeletal muscle^5,7,8^. Yet, whether mitochondrial dynamics are merely downstream consequences of transcriptional programs or are themselves actively instructed by niche-derived metabolic cues remains unclear. Understanding how nutrient availability and redox state integrate with mitochondrial remodeling to guide myogenic progression is an emerging area of interest.

The transitions from quiescence to proliferation and from proliferation to differentiation commitment represent key inflection points in the myogenic program^9^. Both steps require precise metabolic coordination, with growing evidence highlighting mitochondrial function as a central determinant of myogenic progression^10^. Notably, the progression from proliferation to commitment is a developmental milestone that shifts cells from a proliferative progenitor state to an irreversible differentiation fate. This transition might impose a higher threshold for metabolic readiness, requiring the coordinated regulation of nutrient availability, redox balance, and mitochondrial fitness to ensure the successful execution of the myogenic program.

A key regulator of redox homeostasis in this context is glutathione (GSH), the major cellular antioxidant that is maintained through synthesis and recycling under the transcriptional control of Nrf2^11^. During periods of oxidative stress, mitochondrial import of GSH is essential for preserving organelle integrity and preventing oxidative damage^12^. This import is mediated by the mitochondrial GSH carrier Slc25a39, whose expression is dynamically regulated by intracellular GSH levels^13–15^. While Slc25a39 is recognized for its role in iron–sulfur cluster assembly and respiratory chain maintenance, its contribution to redox-sensitive fate transitions in stem cells remains poorly defined^14,16^.

In this study, we identify a glutamine-dependent retrograde redox signaling pathway that serves as a metabolic checkpoint, coupling nutrient availability to myogenic progression through reversible control of mitochondrial dynamics. Under glutamine-limited conditions, proliferating myoblasts orchestrate a reductive redox shift, mitochondrial depolarization, glycolytic suppression, and mitochondrial hyperfusion. These changes, induced by the increased expression of the mitochondrial GSH importer Slc25a39, interrupt differentiation commitment and redirect cells into a reversible, reserve-like state reminiscent of quiescence. Such transient glutamine limitations may naturally occur within the regenerating niche during early phases of capillary rarefaction, before nutrient delivery is restored by infiltrating macrophages and revascularization^17–20^. Thus, this glutamine-dependent redox checkpoint likely functions to preserve stem cell identity under metabolically unfavorable conditions, safeguarding the regenerative process.

## Results

### Active glutamine metabolism sustains glycolysis, mitochondrial fission, and proliferation in myoblasts

Actively proliferating, uncommitted myoblasts utilize glucose and glutamine as primary bioenergetic fuels. To systematically determine the metabolic roles of these nutrients, we isolated MuSCs from hindlimb muscles of wild-type (WT) C57BL/6 mice and cultured them in myoblast growth medium with or without glucose or glutamine for 24 hours. While glucose withdrawal led to profound cell death and cytotoxicity, glutamine depletion resulted in growth arrest without overt toxicity. Under glutamine-replete conditions (+Q), myoblasts remained alive for an extended duration and exhibited circular, donut-shaped mitochondria characteristic of cells in a proliferative state. In contrast, glutamine-deprived (–Q) myoblasts retained an elongated, interconnected mitochondrial network reminiscent of quiescent MuSCs in their physiological niche, despite exposure to mitogenic signals (Figure 1A–B). These findings are consistent with previous studies, which show that active glutamine metabolism is required for myoblast proliferation^19,21,22^. They also suggest a link between extracellular glutamine level and mitochondrial shape dynamics in proliferating myoblasts.

**Figure 1.**
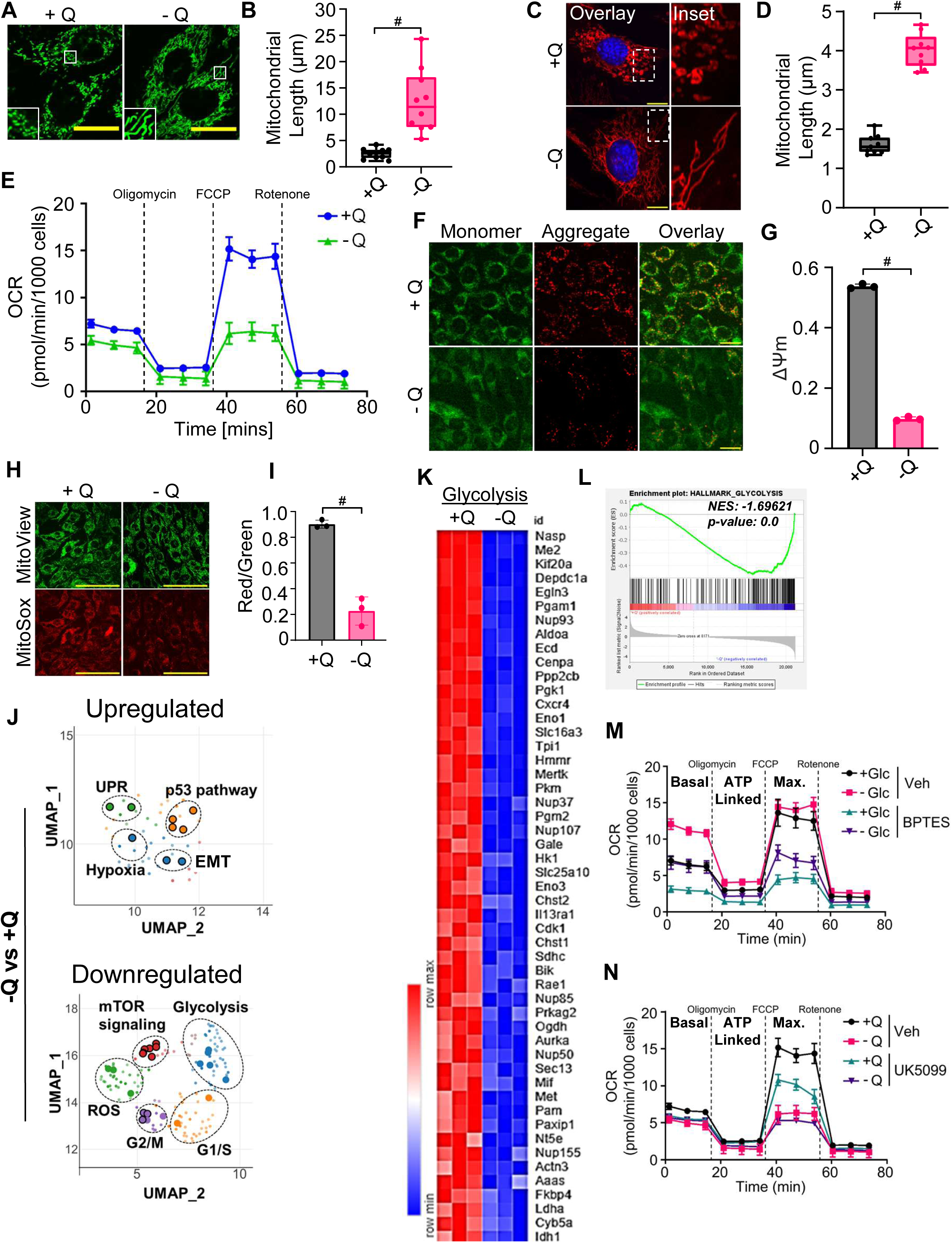
Glutamine deprivation alters mitochondrial morphology and suppresses glycolysis, inducing a Poised Metabolic Arrest (PMA) state in myoblasts. (A) Live-cell imaging of freshly isolated primary myoblasts after culturing in normal growth media with glutamine (+Q) or glutamine-free media (−Q) for 24 hours, stained with MitoView Green to visualize mitochondria. Scale bar = 20 μm. (B) Quantification of mitochondrial length in primary myoblasts under +Q and −Q conditions. Data represent mean ± SD from 10 cells per condition. Statistical significance was determined using Student’s t-test. (C) Immunofluorescence microscopy images of proliferating C2C12 cells stably expressing MitoTag-mCherry cultured in +Q or −Q media, highlighting mitochondrial morphology. Scale bar = 10 μm. (D) Quantification of mitochondrial length in C2C12 cells under +Q and −Q conditions. Data represent mean ± SD from 10 cells per condition/experiment. At least three independent biological replicates were analyzed. Statistical significance was determined using Student’s t-test. (E) Measurement of oxygen consumption rate (OCR) in C2C12 cells cultured in +Q or −Q conditions using the Seahorse XF Analyzer. Data represent mean ± SD from 6 replicates per condition. Statistical analysis was performed using Student’s t-test. (F) Fluorescence microscopy images showing mitochondrial membrane potential (ΔΨm) in proliferating C2C12 cells cultured in +Q or −Q media, assessed using JC-1 dye. Scale bar = 20 μm. (G) Quantification of ΔΨm in C2C12 cells under +Q and −Q conditions. ΔΨm was quantified by calculating the ratio of fluorescence intensity at 488 nm (green monomers) to 568 nm (red aggregates) from JC-1 staining using FIJI software. Data represent mean ± SD from n = 3 independent experiments. Statistical significance was determined using Student’s t-test. (H) Fluorescence microscopy images showing mitochondrial superoxide levels in proliferating C2C12 cells cultured in +Q and −Q conditions, assessed using MitoSOX Red staining. Scale bar = 100 μm. (I) Quantification of mitochondrial superoxide levels in C2C12 cells under +Q and −Q conditions, calculated as the ratio of MitoSOX Red fluorescence (superoxide indicator) to MitoView Green fluorescence (mitochondrial marker). Data represent mean ± SD from n = 3 independent experiments. Statistical analysis was performed using Student’s t-test. (J) Uniform Manifold Approximation and Projection (UMAP) analysis of whole-cell mRNA sequencing data from proliferating C2C12 cells cultured in +Q or −Q media, highlighting upregulated and downregulated gene ontology pathways under glutamine-free conditions. (K) Heatmap displaying the expression levels of the top 51 downregulated glycolysis pathway genes in C2C12 cells under −Q conditions compared to +Q, generated using k-means clustering. (L) Gene Set Enrichment Analysis (GSEA) of glycolysis pathway genes in C2C12 cells cultured in −Q versus +Q conditions, indicating significant downregulation of glycolytic genes upon glutamine deprivation. (M) Measurement of OCR in C2C12 cells cultured in normal growth media containing glucose (+Glc) or glucose-free media (−Glc), treated with BPTES (a glutaminase inhibitor) or vehicle control (DMSO), using the Seahorse XF Analyzer. Data represent mean ± SD from n = 6 replicates per condition. Statistical significance was assessed using Student’s t-test. (N) Measurement of OCR in C2C12 cells cultured in +Q or −Q media, treated with UK5099 (an inhibitor of the mitochondrial pyruvate carrier) or vehicle control (DMSO), using the Seahorse XF Analyzer. Data represent mean ± SD from 6 replicates per condition. Statistical significance was assessed using Student’s t-test. Statistical significance is indicated as *P<0.05(*)* and *P<0.05(#)*, determined by Student’s t-test. Data presented as mean ± SD.

To determine whether the mitochondrial remodeling observed under glutamine deprivation is specific to MuSC activation or reflects a broader feature of myoblast metabolic regulation, we turned to the C2C12 myoblast cell line. C2C12 cells offer a biochemically tractable and well-characterized model that can be maintained in a proliferative state at low confluency and have been extensively used to dissect metabolic fuel flexibility, redox balance, and mitochondrial function^23–25^. We asked whether proliferating C2C12 myoblasts retain the capacity to remodel their mitochondrial networks in response to acute glutamine withdrawal, and whether this response reflects engagement of conserved molecular machinery that links glutamine metabolism to mitochondrial dynamics. Consistent with our findings in primary MuSCs, glutamine deprivation in C2C12 cells stably expressing MitoTag-mCherry induced a significant increase in mitochondrial length and interconnectivity (Figure 1C–D). To determine whether this morphological remodeling is accompanied by changes in mitochondrial function, we next assessed key bioenergetic parameters in glutamine-deprived C2C12 myoblasts. Glutamine withdrawal markedly suppressed mitochondrial oxygen consumption rate (OCR; Figure 1E), consistent with a shift away from oxidative metabolism despite the continued presence of high glucose in the culture media. This was accompanied by a significant reduction in mitochondrial membrane potential (ΔΨm; Figure 1F–G), as well as a decrease in mitochondrial superoxide production (Figure 1H–I), further indicating a globally suppressed mitochondrial activity state.

We next asked whether these metabolic and mitochondrial changes impact myoblast proliferation. Indeed, glutamine-depleted cells showed markedly reduced proliferation and exhibited a marked reduction in BrdU incorporation, indicating impaired S-phase entry (Figure S1A-C). This proliferation arrest is fully reversible, as replenishment of glutamine, even after 3 days of glutamine limitation, results in S-phase initiation without any overt cell death (Figure S1D-E). Together, these data demonstrate that glutamine is required for myoblasts to traverse S-phase, with its absence imposing a reversible proliferation arrest without overt cell death.

To gain a comprehensive understanding of the arrested cell state acquired upon glutamine limitation and identify the molecular pathways leading to increased mitochondrial length, loss of mitochondrial functionality and S-phase arrest under these conditions, we performed whole-cell mRNA sequencing (RNA-seq) analysis on proliferating C2C12 myoblasts cultured in the presence or absence of glutamine. After quality control and alignment, we identified 2128 differentially expressed genes (DEGs) with adjusted p-value (pAdj) <0.05, and |log2 fold change|> 1. Specifically, 513 genes were upregulated, and 1615 genes were downregulated in glutamine-depleted myoblasts. In agreement with previous studies indicating the requirement of glutamine metabolism for myoblast proliferation^21,26,27^ Uniform Manifold Approximation and Projection (UMAP)^28^ analysis of the transcriptomic data under glutamine depletion revealed a significant downregulation of genes associated with mTORC1 signaling and cell cycle progression (G2-M transition) while inducing transcript levels of genes linked to the Unfolded Protein Response (UPR), p53 network, and Epithelial-to-Mesenchymal Transition (EMT) (Figure 1J, Figure S1F).

Together, these transcriptomic and metabolomic changes suggest that glutamine withdrawal reprograms myoblasts into a distinct metabolic state characterized by reduced mitochondrial activity, suppressed glycolysis, and impaired cell cycle progression. Notably, this state shares some features of quiescence and those of the myogenic reserve cell (MRC) population that arises during differentiation. MRCs are a subset of mononucleated myogenic progeny that resist terminal fusion, remain unfused after 48–72 hours under differentiating conditions, and adopt a quiescent-like state. Consistent with prior studies that identify MRCs as molecularly similar to adult quiescent MuSCs, our RNA-seq analysis revealed hallmark features shared by MRCs and quiescence. Thus, we observed reduced *MyoD* transcript levels, accompanied by increased Sprouty1 and Cdkn1a (p21CIP1), along with decreased *MKi67*, and a higher proportion of Pax7-positive myoblasts under glutamine-limited conditions. In contrast to myogenic reserve cells, which arise during differentiation in a mitogen-restrictive environment and upregulate Cdkn1b (p27KIP1), glutamine-limited cells suppressed Cdkn1b, indicating that they remain poised to re-enter the cell cycle. On this basis, we define a reversible, metabolically induced, nonproliferative state termed Poised Metabolic Arrest (PMA). PMA is a unique cellular state entered by proliferating myoblasts in response to glutamine withdrawal, characterized by suppressed mitochondrial function, redox remodeling, and transcriptional adaptation, yet fully capable of reinitiating cell cycle progression when glutamine is restored. Thus, PMA shares features with MRC and quiescent stem cells, but arises from a cell population in response to glutamine limitation.

To further characterize cells in PMA, we analyzed differentially expressed gene (DEG) signatures (Figure 1K), and Gene Set Enrichment analysis, which revealed that glutamine depletion markedly repressed genes involved in glycolysis (Figure 1L, NES: −1.69621, P-value: 0.0). mRNA levels of important glycolytic genes such as *Pgk1 (*pAdj= 0.0012), *Pgm2* (pAdj= 0.0029), *Slc16a3* (pAdj= 0.002), *Me2* (pAdj= 0.0006), and others were lower in glutamine-depleted cells (−Q) compared with control (+Q) (Figure 1K-L). Furthermore, unbiased metabolomic analysis revealed a significant reduction in levels of glycolytic intermediates under glutamine-depleted conditions (Figure S1G). Taken together, the evidence suggests that PMA cells exhibit a repressed transcriptional program of glycolysis due to the loss of active glutamine metabolism.

To further confirm this idea, we first examined mitochondrial function when glucose versus glutamine limitation. Under glucose limitation, myoblasts showed increased basal and ATP-linked oxygen consumption rate. This increase reflected a compensatory reliance on oxidative phosphorylation through glutaminolysis, as evidenced by the glutaminase inhibitor BPTES, which reduced both basal and ATP-linked oxygen consumption, and also decreased the extracellular acidification rate under glucose limitation (Figures 1M and S1H). Despite this compensation, glucose withdrawal lowered maximal uncoupled respiration, indicating that glucose contributes to spare respiratory capacity in proliferating myoblasts. Consistent with that interpretation, the inhibition of the mitochondrial pyruvate carrier 1 (MPC1) with UK5099 did not change basal or ATP-linked respiration but markedly reduced maximal respiration in standard glutamine-replete medium.

Under glutamine-depleted conditions, basal, ATP-linked, and maximal oxygen consumption were all suppressed, and inhibition of the mitochondrial pyruvate carrier had no additional effect (Figure 1N). Glutamine depletion also caused a near-complete drop in extracellular acidification rate, a proxy for glycolysis (Figure S1I). Given the very low oxygen consumption in the absence of glutamine, the reduction in extracellular acidification is best explained by impaired glycolytic flux rather than increased pyruvate delivery into the mitochondria. This view is supported by two observations. First, the inhibition of the MPC1 did not lower ATP-linked respiration under either glutamine state. Second, unbiased metabolomics showed decreased levels of glycolytic intermediates under glutamine depletion (Figure S1G). Thus, we conclude that in proliferating myoblasts, glycolysis contributes to maximal respiration when glutamine metabolism is active.

Taken together, these findings suggest that the necessity of active glutamine metabolism in myoblasts is to support both glycolysis and oxidative phosphorylation, serving as a central hub that links mitochondrial bioenergetics with the transcriptional control of proliferation and differentiation. In its absence, cells respond by entering a reversible PMA state, characterized by the suppression of glycolysis, mitochondrial remodeling, and the reprogramming of cell cycle and fate determinants, and remain poised to regain full differentiation capacity upon glutamine restoration.

### Increased cellular GSH levels contribute to mitochondrial remodeling in glutamine-depleted conditions

To investigate how glutamine metabolism influences mitochondrial length and metabolic features of the PMA state, we examined the functional roles of its downstream metabolic intermediates. Beyond ATP production, mitochondrial glutamine oxidation redistributes glutamine-derived carbon and nitrogen across key metabolic pathways^29^. For example, glutamine-derived alpha-ketoglutarate (α-KG) is a primary branching point where glutamine carbons can be channeled toward GSH biosynthesis, epigenetic regulation, and the TCA cycle (Figure 2A). Consistent with this, adding cell-permeable α-KG nearly completely restored circular mitochondrial morphology in glutamine-depleted cells (Figure 2B-C). Similarly, while succinate supplementation partially reversed mitochondrial fragmentation, only cell-permeable α-KG fully rescued the myogenic impairment due to glutamine withdrawal (Figure S2A-B). These findings indicate that the metabolic fate of glutamine downstream of α-KG is crucial for maintaining mitochondrial morphology and supporting myoblast function.

**Figure 2.**
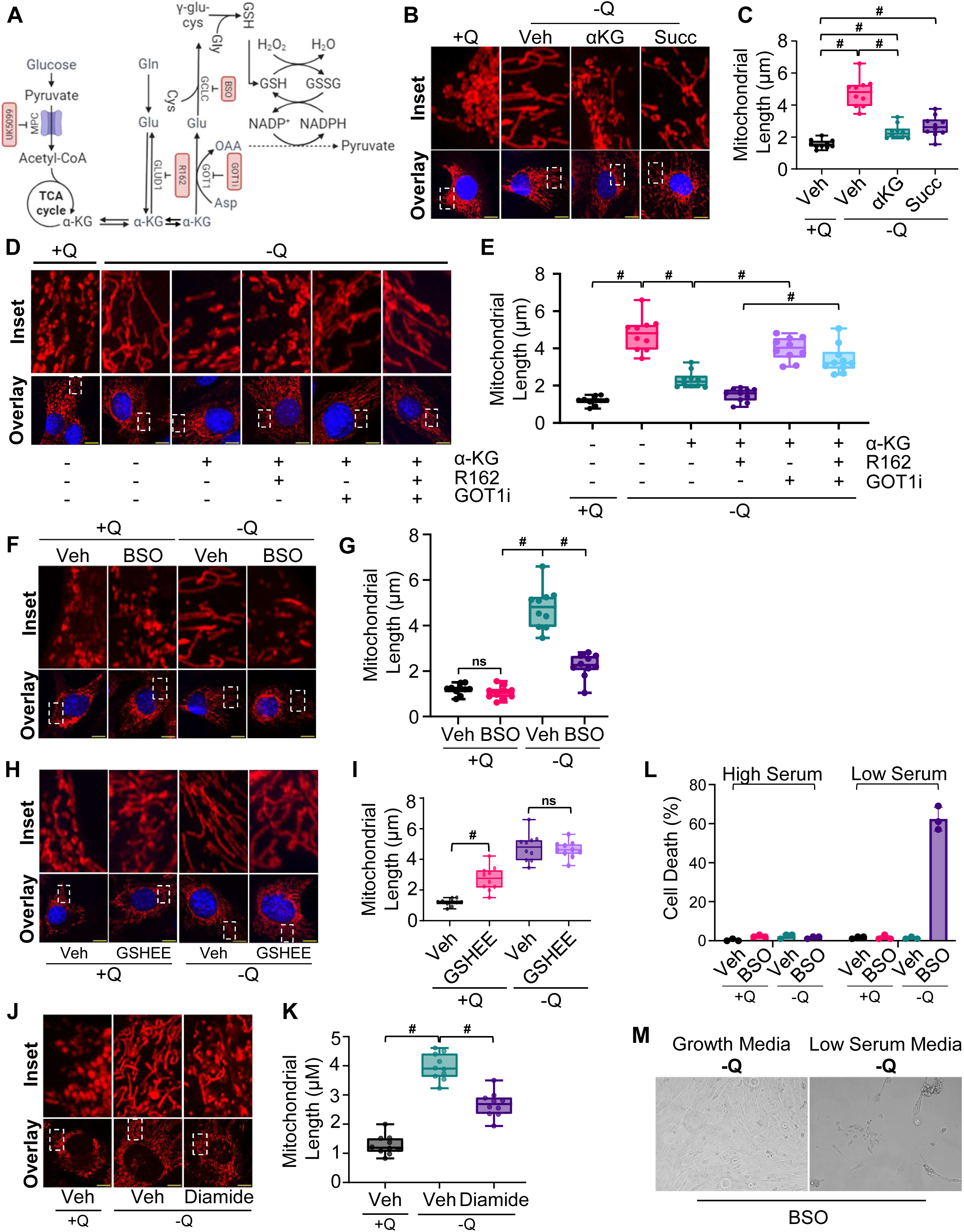
Increased glutathione synthesis is necessary to remodel mitochondria in the absence of glutamine metabolism. (A) Schematic diagram illustrating the metabolic pathways of glutamine-derived carbons, including their roles in glutathione (GSH) biosynthesis, epigenetic regulation, and the tricarboxylic acid (TCA) cycle. (B) Immunofluorescence images of C2C12 MitoTag-mCherry cells cultured in +Q or −Q media, treated with dimethyl α-ketoglutarate (αKG), sodium succinate (Succ), or vehicle control (Veh). Scale bar = 10 μm. (C) Quantification of mitochondrial length under the indicated conditions. Data represent mean ± SD from 10 cells per condition. Statistical significance was determined using Student’s t-test. (D) Immunofluorescence images of cells treated with αKG, R162 (a Glud1 inhibitor), GOT1i (an inhibitor of glutamic-oxaloacetic transaminase 1), or their combinations in +Q or −Q media. Scale bar = 10 μm. (E) Quantification of mitochondrial length under the indicated treatment conditions. Data represent mean ± SD from 10 cells per condition. Statistical significance was determined using Student’s t-test. (F) Immunofluorescence images of cells treated with Buthionine Sulfoximine (BSO, a GSH synthesis inhibitor) or Veh in +Q or −Q media. Scale bar = 10 μm. (G) Quantification of mitochondrial length under the indicated conditions. Data represent mean ± SD from 10 cells per condition. Statistical significance was determined using Student’s t-test. (H) Immunofluorescence images of cells treated with glutathione reduced ethyl ester (GSHEE, a cell-permeable form of GSH) or Veh in +Q or −Q media. Scale bar = 10 μm. (I) Quantification of mitochondrial length under the indicated conditions. Data represent mean ± SD from 10 cells per condition. Statistical significance was determined using Student’s t-test. (J) Immunofluorescence images of cells treated with diamide (a thiol-oxidizing agent) or Veh in +Q or −Q media. Scale bar = 10 μm. (K) Quantification of mitochondrial length under the indicated conditions. Data represent mean ± SD from 10 cells per condition. Statistical significance was determined using Student’s t-test. (L) Cell death measured by Trypan Blue exclusion in cells cultured in glutamine-replete (+Q) or glutamine-free (–Q) media (Day 0) with BSO or vehicle (Veh), compared to cells in low-serum media (Day 2). Data represent mean ± S.D. from four replicates per condition. Statistical significance was determined by Student’s t-test. (M) Brightfield images of cells treated with BSO in glutamine-free growth media (10% FBS) or glutamine-free low-serum media (2% horse serum). Statistical significance is indicated as P < 0.05 (*, #), determined by Student’s t-test. Data are shown as mean ± S.D.

In addition to replenishing the TCA cycle and serving as a co-factor in DNA and histone demethylation reactions, alpha-KG is an important precursor for many biosynthetic reactions in the mitochondria and the cytosol. For example, alpha-KG is transaminated to generate glutamate via the enzyme GOT1, a reaction that also generates oxaloacetate (OAA) from aspartate (Figure 2A)^25,30–32^. A two-step reaction from OAA to pyruvate links glutamine metabolism with glutathione regeneration via the production of NADPH^25,30–32^. Finally, under glutamine-depleted conditions, alpha-KG is also a substrate in a reactive reamination reaction catalyzed by the enzyme Glud1 to generate glutamate^19,27,29^. Thus, due to the multi-directional fates of alpha-KG, we undertook a chemical/genetic analysis to identify the alpha-KG-dependent mechanisms regulating mitochondrial length under glutamine-starved conditions in myoblasts.

We first tested if either transamination of alpha-KG via Got1 or its reactive reamination via Glud1 mediated rescue of mitochondrial length under glutamine-depleted conditions (Figure 2A). Whereas Got1 inhibition by Got1i treatment blunted the effectiveness of alpha-KG in reversing the elongation of mitochondrial length under glutamine-depleted conditions, the inhibition of Glud1 by R162 treatment in alpha-KG-treated cells resulted in a near-complete rescue of mitochondrial length under glutamine-depleted conditions (Figure 2D-E). Furthermore, Got1, but not Glud1 inhibition, prevented the rescue of myogenesis by alpha-KG under glutamine-depleted conditions (Figure S2C-D). These results indicate that metabolic pathways downstream of glutamine, involving Got1, play a crucial role in regulating mitochondrial length and myogenesis in myoblasts, and that the near-complete rescue of mitochondrial length in Glud1-inhibited cells is likely due to the diversion of alpha-KG to the Got1 enzymatic pathway.

Got1 is a key cytosolic enzyme downstream of glutamine that diversifies the fates of alpha-KG towards glutathione (GSH) regeneration and amino acid biosynthesis pathways^26,27^. Therefore, wondered if glutamine regulation of mitochondrial length is mediated through alterations in GSH metabolism. GCL is a rate-limiting enzyme for GSH biosynthesis and can be inhibited by buthionine sulfoximine (BSO), which decreases intracellular GSH levels^33,34^ (Figure 2A). Treatment with BSO significantly attenuated the increases in mitochondrial length observed in glutamine-deprived cells without affecting control cells (Figure 2F-G). On the other hand, increasing the intracellular pool of GSH with cell-permeable GSH ethyl ester (GSHEE) markedly increased mitochondrial length in myoblasts cultured in the presence of glutamine without affecting those cultured in the absence of glutamine (Figure 2H-I). Conversely, diamide treatment, which converts oxidized glutathione (GSH) to reduced glutathione (GSSG), modestly but significantly reversed the mitochondrial length of glutamine-depleted cells (Figure 2J-K). These data strongly suggest that an increased ratio of GSH:GSSG in glutamine-depleted cells drives the elongation of mitochondrial length. Consistently, glutamine-depletion of myoblasts increased whole-cell reduced GSH levels (Figure S2E). Elevated intracellular GSH levels, decreased oxidative phosphorylation capacity, reduced mitochondrial ROS, and membrane potential are some of the hallmarks of cellular reductive stress^35–37^. Therefore, we conclude that glutamine-deprived cells experience reductive stress, leading to the establishment of features similar to mitochondrial respiratory quiescence (MRQ)^38,39^.

### Increased cellular GSH levels are cytoprotective in glutamine-starved myoblasts

Unlike many normally proliferative cell types, such as cancer cells or immortalized fibroblasts, myoblasts are resistant to the cytotoxicity associated with glutamine depletion and remain viable in culture media for an extended duration, despite not proliferating^21,32^. To test whether the cellular reductive environment provides cytoprotective features, we inhibited GSH synthesis by treating cells with BSO in the presence or absence of glutamine under low (2% v/v) or high (10% v/v) serum conditions. While BSO treatment was well-tolerated by cells cultured in the presence or absence of glutamine at high serum concentrations, glutamine-depleted cells strikingly failed to survive low serum conditions when GSH synthesis was inhibited by BSO treatment, indicating that inhibition of GSH biosynthesis in the absence of pro-survival inputs from serum sensitizes glutamine-depleted myoblasts, resulting in profound cell death (Figure 2L-M). Taken together, our characterization of myoblasts cultured in glutamine-deficient media suggests a distinctive metabolic adaptation to glutamine withdrawal. Under these conditions, myoblasts repress glycolytic gene transcription, induce mitochondrial elongation and mitochondrial respiratory quiescence, and modify GSH metabolism. These adaptations protect against cytotoxicity (Figure 2L-M), pause the cell cycle progression (Figure S1A-C), and prevent differentiation (Figure S2A-D)^21^.

### Selective solute carrier (Slc) transcriptome rewiring supports GSH biosynthesis under glutamine-depleted conditions

In the absence of active mitochondria in glutamine-depleted myoblasts, the NADPH-dependent glutathione regeneration circuit is unlikely to function efficiently^29,32^. Yet, we observed elevated intracellular reduced GSH levels, indicating that myoblasts adapt to glutamine depletion by altering either GSH utilization or the GSH biosynthetic pathway. To identify molecular circuits in myoblasts that maintain elevated GSH levels in glutamine-depleted cells, we performed KEGG metabolic pathway analysis of differentially expressed transcripts under glutamine-depleted conditions. Consistent with our determination that glutamine depletion leads to quiescent MuSCs-like features, we observed increased expression of genes driving fatty acid oxidation (FAO) (Figure 3A), a primary fuel in quiescent MuSCs. Furthermore, under glutamine-depleted (−Q) conditions, we also observed increases in the mRNA levels of genes associated with metabolite importers, autophagy regulators, the unfolded protein response, and amino acid transaminases (Figure 3A). Of these, we noticed a cluster of nutrient and metabolite carriers belonging to the Solute Carrier (SLC) family of transporters that are significantly upregulated in −Q conditions (Figure 3A-C). Solute carriers (Slc) are critical gatekeepers of nutrient flux into and out of cells and organelles. Slc family members acutely respond to and compensate for nutrient deficiencies, imparting metabolic plasticity to the cells^40^. Our analyses identified the upregulation of several classes of plasma membrane amino acid transporters, including neutral amino acids (Slc1a4), cysteine (Slc7a11, Slc3a2), glutamate (Slc1a3), arginine (Slc7a3), glycine (Slc6a9), and aspartate (Slc1a4) (Figure 3C-D). Glutamine depletion also significantly downregulated the plasma membrane urea transporter Slc14a1, potentially due to a decrease in glutamine cleavage-derived ammonia (Figure 3C-D). In addition, several members of the mitochondrial solute carrier family were upregulated in glutamine-depleted conditions. Among these, Slc25a39, which encodes a recently identified mitochondrial glutathione importer^15,41^, was the most significantly upregulated mRNA amongst the mitochondrial Slc25 family of transporters in glutamine-depleted myoblasts (Figure 3E).

**Figure 3.**
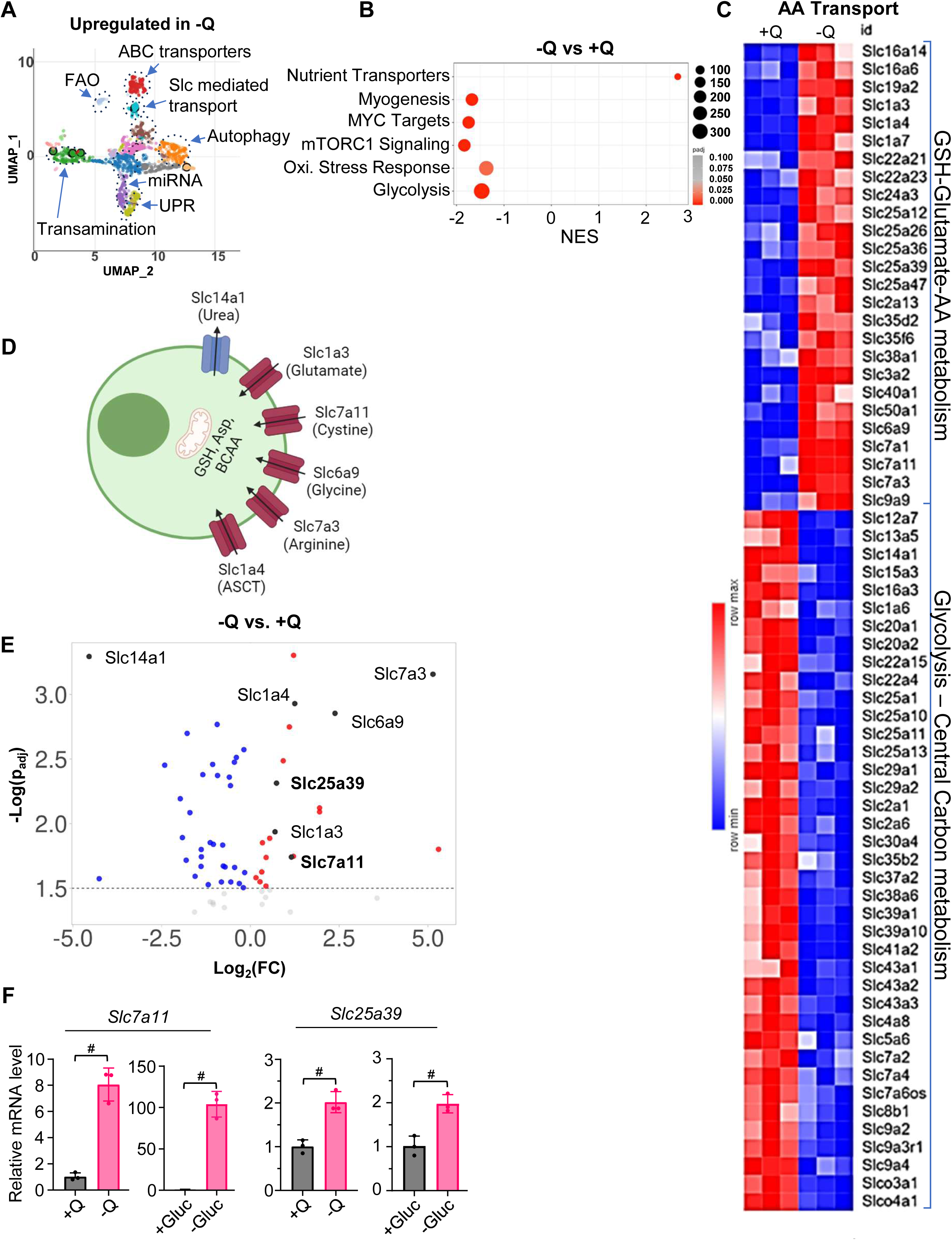
Solute carrier transcriptome remodeling underscores increased GSH biosynthesis under glutamine limitation. (A) UMAP analysis of KEGG pathways upregulated in C2C12 cells cultured under −Q conditions compared to +Q. (B) Cluster plot from GSEA comparing gene expression profiles of cells in −Q versus +Q media, highlighting significantly enriched gene sets. (C) Heatmap displaying expression levels of amino acid transporter genes under −Q and +Q conditions, generated using k-means clustering. (D) Highlighted expression levels of specific solute carrier (SLC) transcripts involved in glutathione synthesis, including *Slc7a11* and *Slc25a39*, under −Q and +Q conditions. (E) Volcano plot showing differential expression of SLC gene family mRNAs under −Q conditions compared to +Q, indicating upregulated and downregulated genes. (F) qRT-PCR analysis of *Slc7a11* and *Slc25a39* mRNA expression levels in cells cultured under −Q or glucose-free (−Gluc) conditions. Data represent mean ± SD from n = 3 independent experiments. Statistical significance was determined using Student’s t-test. Statistical significance is indicated as *P<0.05(*)* and *P<0.05(#)*. Data presented as mean ± SD.

As the transport activities of Slc25a39 and Slc7a11 are involved in GSH metabolism^14,42–45^, we further characterized the transcriptional regulation and function of these transporters in myoblasts. We first validated the upregulation of Slc7a11 and Slc25a39 mRNA levels using qRT-PCR under glutamine- or glucose-depleted conditions. Consistent with previous reports, the mRNA level of Slc7a11 was significantly elevated in myoblasts in response to glucose withdrawal^46^. We also observed a significant increase in Slc7a11 mRNA levels after glutamine withdrawal (Figure 3F). In addition, the mRNA levels of Slc25a39 were markedly increased upon glutamine and glucose starvation (Figure 3F). These findings validate RNAseq results, confirming that Slc7a11 and Slc25a39 are transcriptional targets of the pathway activated under nutrient-starved conditions in myoblasts.

To identify the upstream transcription factor network that regulates Slc25a39 expression under glutamine-depleted conditions, we queried the transcription factor databases with the top 250 significantly upregulated transcripts (adj. p<0.05) in glutamine-depleted conditions. We found two clusters of Atf4 and Nrf2 targets that were up-regulated under glutamine-depleted conditions. Recently, Atf4 was identified as an upstream regulator of the Nrf2 transcriptional activity through Chac1-dependent and independent pathways^47–49^. Consistently, we observed a marked increase in Atf3, Atf4, and Chac1 levels under glutamine-depleted conditions in our RNA-seq analysis (Figure S3A). Nrf2 is a downstream target of Atf4 and a major transcriptional regulator of GSH metabolism in cells. In our RNA-seq analysis, we detected upregulation of several Nrf2 target genes, such as *Slc7a11*, *Sqstm1*, and *Abcc4,* under glutamine-depleted conditions (Figure 4A-B). To confirm Nrf2 pathway activation under glutamine-depleted conditions, we quantified the mRNA levels of key regulators and targets of Nrf2 by qRT-PCR analysis. We observed a steep decrease in Nrf2 inhibitor Keap1 (Figure 4A) mRNA levels and an increase in Nrf2 mRNA levels, along with the upregulation of several of the Nrf2 target genes, including *Slc7a11*, *Gclc*, and *Hmox1* (Figure 4C). Consistent with the observation that glutamine deprivation suppresses glycolytic gene expression in C2C12 myoblasts, we found that *Mpc1* mRNA levels were reduced in glutamine-depleted conditions (Figure 4B). Notably, this suppression was reversed upon Nrf2 silencing, suggesting a regulatory interaction between the Nrf2 signaling-GSH biosynthesis axis and the transcriptional control of glycolysis in myoblasts under glutamine deprivation. Glutamine depletion also led to increased mRNA levels of Slc7a11 and Slc25a39, which was abolished when Nrf2 was silenced, indicating that the induction of Slc25a39 operates downstream of Nrf2, likely as a direct transcriptional target, under steady-state and glutamine-starved conditions (Figure 4C).

**Figure 4.**
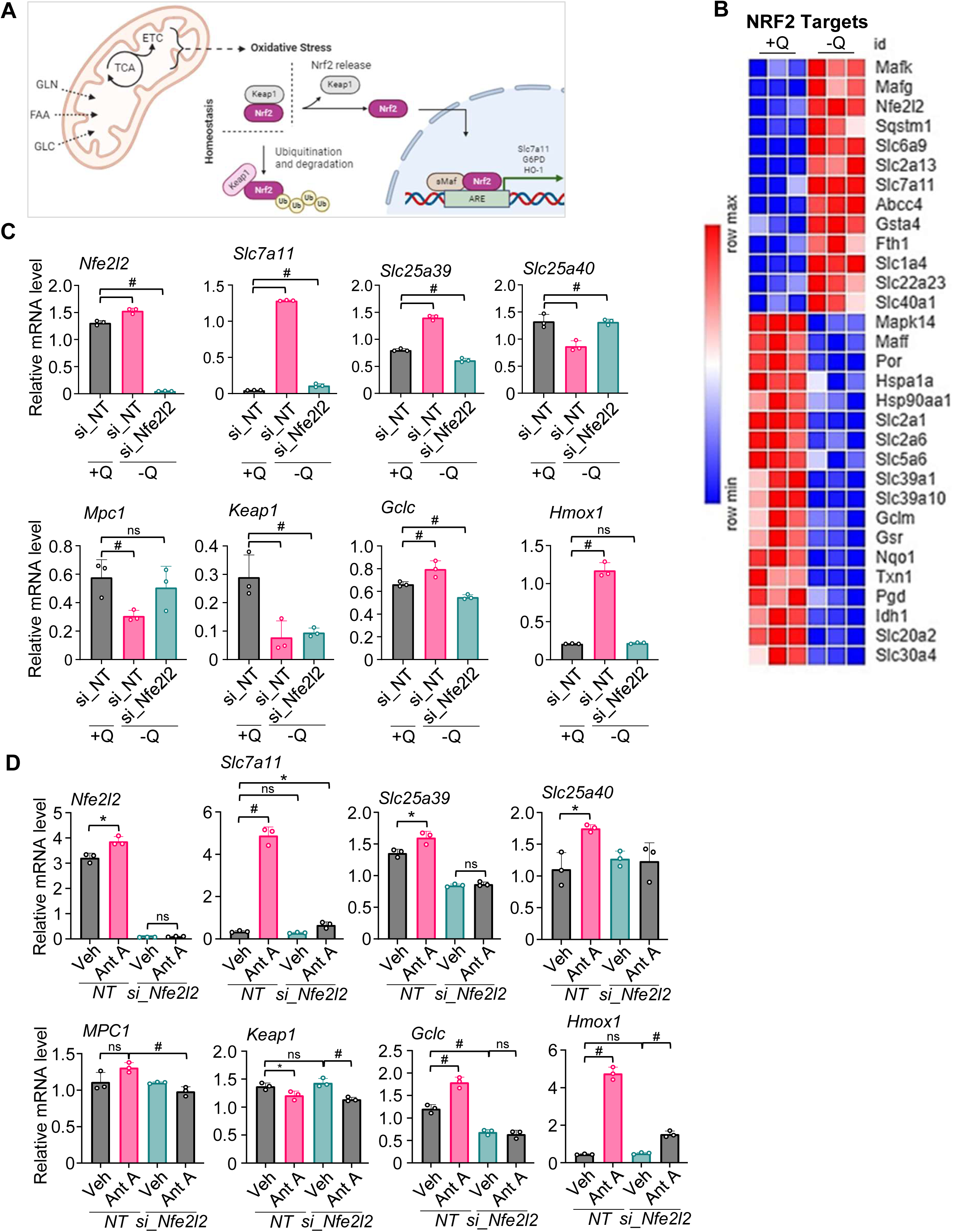
Slc25a39 is a downstream target of the Atf4/Nrf2 network under nutrient or redox stress conditions. (A) Schematic diagram illustrating how metabolic pathways contribute to increased oxidative stress, leading to Nrf2 stabilization and upregulation of the oxidative stress response. (B) Heatmap displaying expression levels of Nrf2 target genes under +Q and −Q conditions, generated using k-means clustering. (C) qRT-PCR analysis of indicated mRNA levels from cells cultured in +Q or −Q media, transfected with non-targeting siRNA (si_NT) or *Nfe2l2* siRNA (si_Nfe2l2). Data represent mean ± SD from n = 3 independent experiments. Statistical significance was determined using Student’s t-test. (D) qRT-PCR analysis of the same genes in cells transfected with si_NT or si_Nfe2l2 and treated with 6.25μM antimycin A (Ant A) or Veh. Data represent mean ± SD from n = 3 independent experiments. Statistical significance was determined using Student’s t-test. Statistical significance is indicated as *P<0.05(*)* and *P<0.05(#)*, determined by Student’s t-test. Data presented as mean ± SD.

Inhibition of mitochondrial Complex III by Antimycin A (Ant A) produces superoxide, triggering cellular oxidative stress that promotes Nrf2 nuclear translocation and activation of Nrf2-dependent gene expression (Figure 4A)^42,50^. This is a physiologically relevant modality that affects cellular redox homeostasis. To determine whether Slc25a39 is a downstream target of the oxidative stress-induced Nrf2 transcriptional network, we treated control and Nrf2-silenced myoblasts with Antimycin A and measured the mRNA levels of the validated Nrf2 downstream targets. As expected, Antimycin A treatment induced the expression of several Nrf2 targets, including *Slc7a11, Gclc,* and *Hmox1*. Antimycin A treatment also significantly induced the expression of *Slc25a39* in myoblasts, which was blunted by Nrf2 silencing (Figure 4D). Thus, *Slc25a39* is a transcriptional target of Nrf2, whose expression is induced under glutamine depletion and oxidative stress in myoblasts.

### Mitochondrial GSH pool regulates mitochondrial dynamics and activity under glutamine-depleted conditions

Glutamine depletion and the resultant increase in cellular GSH biosynthetic pathways remodel mitochondrial length. Given that glutamine depletion upregulated the expression of several mitochondrial transcripts involved in GSH biosynthesis or mitochondrial import, we set out to determine if Slc7a11 and Slc25a39 regulate mitochondrial length under glutamine-depleted conditions. As expected, under glutamine-depleted conditions, the siRNA-mediated silencing of Slc7a11 decreased mitochondrial length, likely due to a reduced intracellular GSH pool. Interestingly, loss of mitochondrial GSH import via Slc25a39 knockdown was equally effective at reducing mitochondrial length in glutamine-depleted cells without affecting mitochondrial length in glutamine-replete cells (Figure 5A-B). This finding suggests that mitochondrial thiol metabolism may be an important determinant of mitochondrial length under glutamine-depleted conditions. To test this hypothesis, we treated glutamine-depleted cells with mitoCDNB, a mitochondrially targeted chlorodinitrobenzene that depletes the mitochondrial GSH pool^37^. While mitoCDNB did not further affect the mitochondrial length in myoblast cells cultured in glutamine-containing media, mitoCDNB treatment increased mitochondrial fragmentation in glutamine-depleted cells, confirming that the mitochondrial thiol balance is a critical determinant of mitochondrial length regulation in the absence of glutamine (Figure 5C-D).

**Figure 5.**
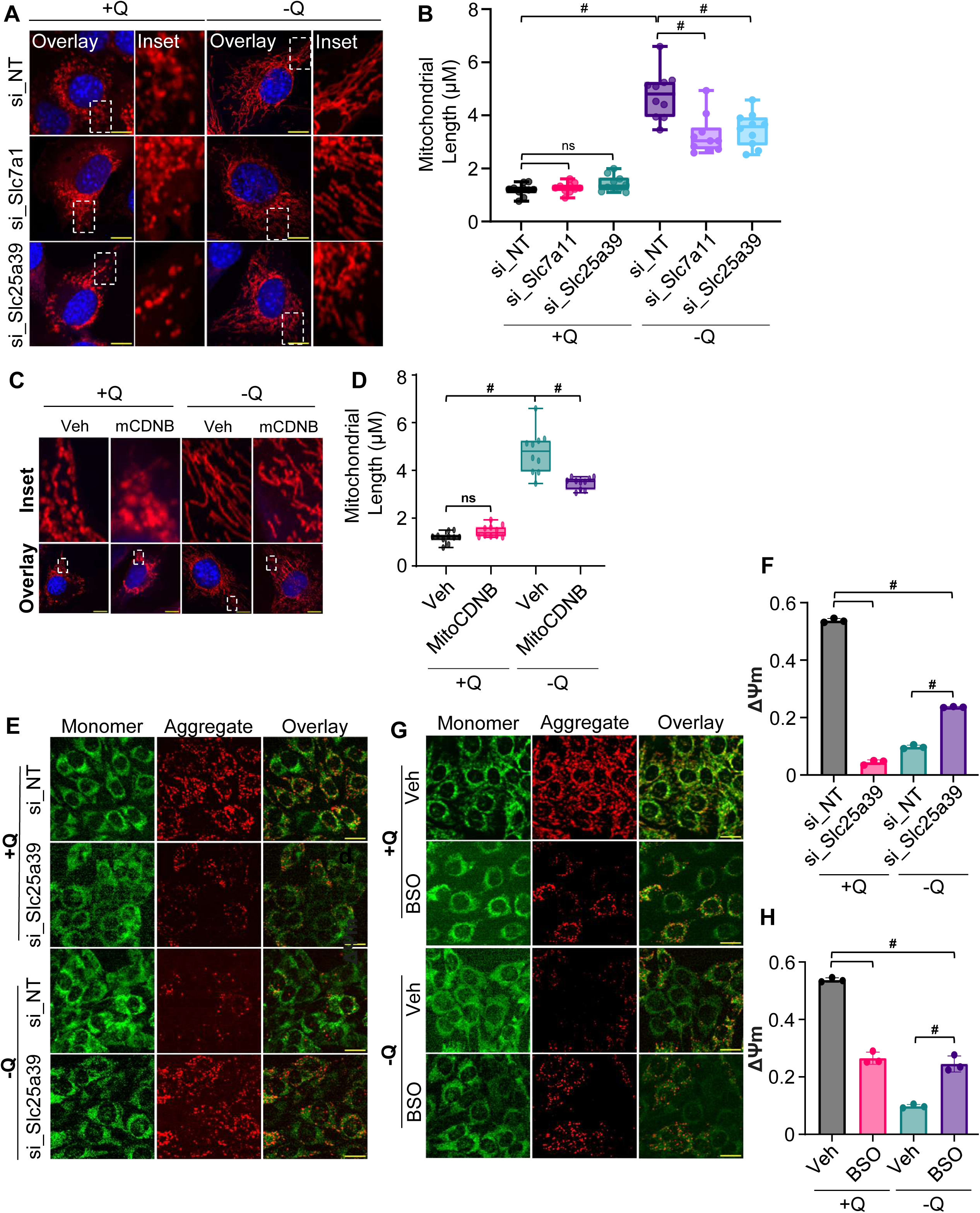
Mitochondrial GSH levels regulate mitochondrial functionality and dynamics under glutamine-depleted conditions. (A) Immunofluorescence microscopy images of C2C12 MitoTag-mCherry cells cultured in +Q or −Q media, transfected with si_NT, *Slc7a11* siRNA (si_Slc7a11), or *Slc25a39* siRNA (si_Slc25a39). Scale bar = 10 μm. (B) Quantification of mitochondrial length under the indicated conditions. Data represent mean ± SD from n = 10 cells per condition. Statistical significance was determined using Student’s t-test. (C) Immunofluorescence images of cells treated with MitoCDNB (mCDNB, a mitochondria-targeted glutathione-depleting agent) or Veh in +Q or −Q media. Scale bar = 10 μm. (D) Quantification of mitochondrial length under the indicated conditions. Data represent mean ± SD from n = 10 cells per condition. Statistical significance was determined using Student’s t-test. (E) JC-1 assay showing ΔΨm in cells cultured in +Q or −Q media, transfected with si_NT or si_Slc25a39. Scale bar = 20 μm. (F) Quantification of ΔΨm calculated as the ratio of JC-1 fluorescence intensity at 488 nm to 568 nm using FIJI software. Data represent mean ± SD from n = 3 independent experiments. Statistical significance was determined using Student’s t-test. (G) JC-1 assay showing ΔΨm in cells treated with BSO or Veh in +Q or −Q media. Scale bar = 20 μm. (H) Quantification of ΔΨm under the indicated conditions. Data represent mean ± SD from n = 3 independent experiments. Statistical significance was determined using Student’s t-test. Statistical significance is indicated as *P<0.05(*)* and *P<0.05(#)*, determined by Student’s t-test. Data presented as mean ± SD.

Glutamine is a primary fuel for proliferating myoblasts, supporting oxidative phosphorylation (Figure 1E) and the maintenance of mitochondrial membrane potential (Figure 1F). Given that Slc25a39 expression is induced under glutamine depletion or oxidative stress conditions in cultured myoblasts (Figure 3F, Figure 4C-D) and that Slc25a39 silencing partially rescues mitochondrial length under glutamine-depleted conditions (Figure 5A-B), we asked whether Slc25a39 silencing affects mitochondrial functionality under glutamine-depleted conditions. As a measure of mitochondrial function, we quantified the effect of Slc25a39 loss on the mitochondrial membrane potential in myoblasts cultured in the presence or absence of glutamine. Consistent with its role in regulating mitochondrial thiol metabolism, the loss of Slc25a39 in the glutamine-replete condition resulted in a marked loss of mitochondrial membrane potential, as indicated by a decrease in the ratio between oligomeric and monomeric forms of JC-1 (Figure 5E-F). Unexpectedly, when combined with glutamine depletion, the loss of Slc25a39 partially restored mitochondrial membrane potential (Figure 5E-F). Since glutamine depletion induces reductive stress in myoblasts, these results suggest that silencing Slc25a39 could partially mitigate mitochondrial respiratory quiescence under glutamine-depleted conditions, reactivating mitochondrial function.

Thus, we propose that glutamine levels and Slc25a39 expression coordinate mitochondrial membrane potential by acutely regulating mitochondrial oxidative/reductive stress balance. Consistent with this notion, inhibiting total GSH biosynthesis using BSO decreased membrane potential in glutamine-replete cells. In contrast, the same treatment rescued some of the lost membrane potential under glutamine-depleted conditions (Figure 5G-H). Thus, precise coordination between mitochondrial glutamine metabolism and Slc25a39 levels is necessary to maintain the optimal mitochondrial function in proliferating myoblasts.

### Slc25a39 links GSH metabolism to glycolysis repression in glutamine-depleted cells via feedback modulation of Nrf2 activity

To elucidate the transcriptional adaptations underlying improved mitochondrial membrane potential in response to combined conditions of Slc25a39 knockdown and glutamine depletion, we conducted RNA-seq analysis across Control (+Q), glutamine depletion (−Q), Slc25a39 knockdown (KD), and the combined treatment of glutamine depletion with Slc25a39 knockdown(−QKD) conditions. Using a statistically significant differential expression threshold (|log2 fold change| ≥ 1, pAdj ≤ 0.05), we identified ∼900 significantly upregulated genes across all conditions (Figure 6A, Table S1). Notably, the −QKD condition showed the most significant transcriptional response, with 594 genes (70% of total DEG under −QKD conditions) upregulated relative to Control, compared to 242 (16% of total DEG under −Q conditions) genes in −Q and 54 (8% of total DEG under KD conditions) genes in KD, indicating a pronounced transcriptional response in −QKD relative to either treatment alone (Table S1).

**Figure 6.**
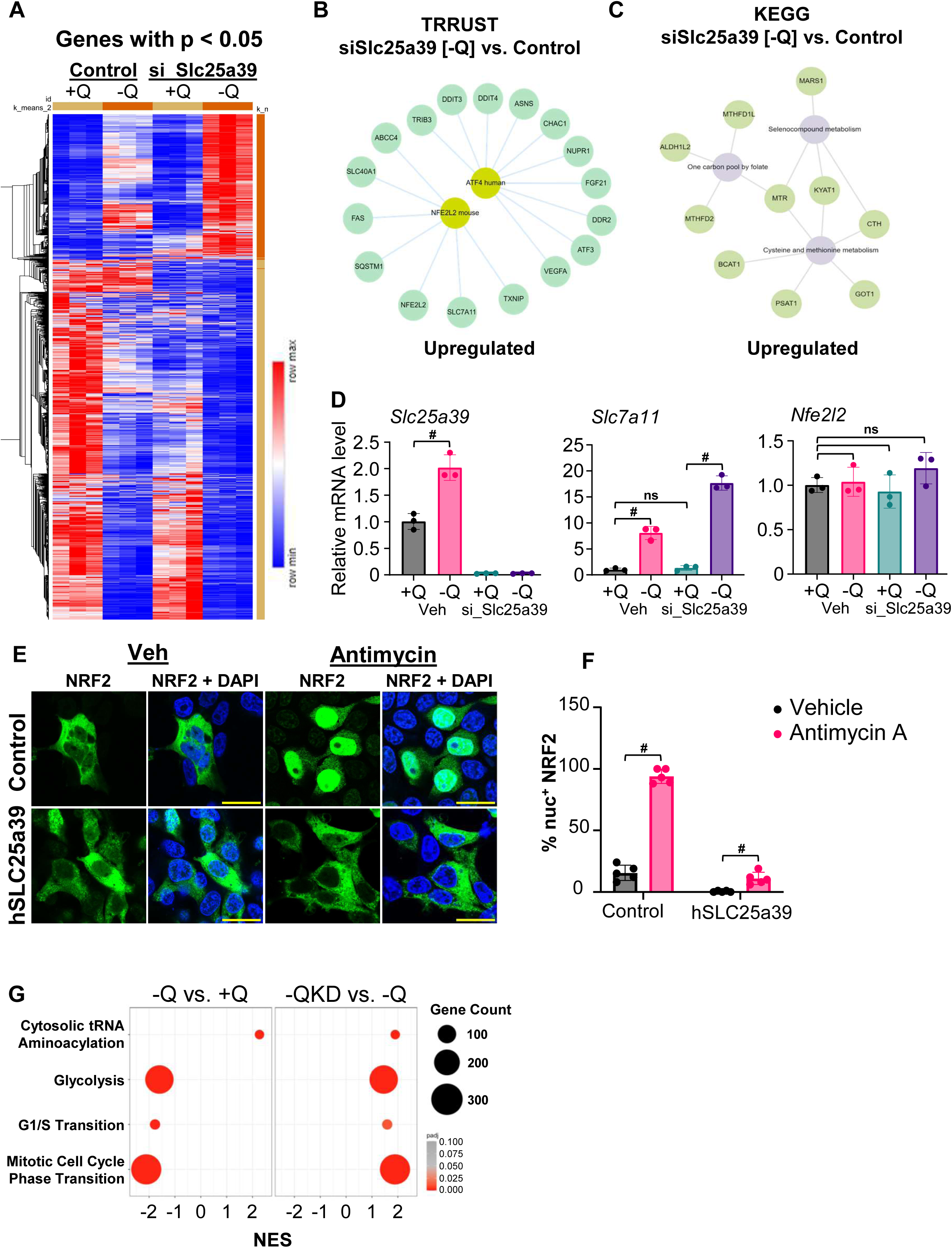
Slc25a39 is necessary for the establishment of the PMA state under glutamine limitations via feedback regulation of Nrf2. (A) Heatmap displaying gene expression profiles (k-means clustering) of cells under four conditions: control in +Q media (Control, +Q), control in −Q media (Control, −Q), *Slc25a39* siRNA knockdown in +Q media (si_Slc25a39, +Q), and *Slc25a39* siRNA knockdown in −Q media (si_Slc25a39, −Q). (B) Top pathways upregulated in glutamine-free *Slc25a39* siRNA knockdown (−QKD) compared to control, identified using the TRRUST library via Enrichr-KG analysis. (C) Top pathways upregulated in −QKD compared to control, identified using the KEGG 2021 library via Enrichr-KG analysis. (D) qRT-PCR validation of *Slc25a39*, *Slc7a11*, and *Nfe2l2* mRNA expression levels in cells cultured in +Q or −Q media, with or without *Slc25a39* siRNA knockdown. Data represent mean ± SD from n = 3 independent experiments. Statistical significance was determined using Student’s t-test. (E) Immunofluorescence microscopy images of WT C2C12 cells and inducible SLC25A39 overexpression cells (iSLC25A39) cultured in +Q media, treated with antimycin A or Veh (DMSO). Nrf2 nuclear localization was assessed. Scale bar = 20 μm. (F) Quantification of Nrf2 nuclear localization. Data represent mean ± SD from n = 5 fields per condition. Statistical significance was determined using Student’s t-test. (G) Cluster plot analysis comparing −Q to +Q conditions (left) and −QKD to −Q conditions (right), highlighting differentially expressed genes and pathway enrichments. Statistical significance is indicated as *p*< 0.05 and *P<0.05(#)*, determined by Student’s t-test. Data are presented as mean ± S.D.

To functionally bin these transcriptional changes across conditions, we applied hierarchical *k*-means clustering and principal component analysis (PCA) (Figure S4A), which together revealed distinct patterns of gene regulation (Figure S4B) that co-dependent upon glutamine availability and Slc25a39 expression. Thus, PCA indicated that −QKD samples exhibited the most distinctive profile, clustering separately along all three components (PC1, PC2, and PC3) compared to individual treatment groups. This pattern reflects a unique transcriptional response in −QKD conditions when both glutamine and Slc25a39 are absent (Figure S4A). Furthermore, *k-*means clustering identified five distinct clusters, each representing transcriptional patterns associated with glutamine metabolism, Slc25a39 expression levels, and the combined effects of both (Table S2, Figure S4B). Notably, clusters 1-2 comprised 381 out of 572 upregulated clustered genes (∼66% of all upregulated genes) whose expression remained unaffected by Slc25a39 knockdown in the presence of glutamine and showed only modest increases under glutamine-depleted (−Q) conditions. However, the combined loss of glutamine and Slc25a39 (−QKD) resulted in a dramatic induction of their expression, suggesting an epistatic interaction between Slc25a39 function and glutamine metabolism, regulating these gene sets (Figure S4B). This distinct clustering pattern observed in the −QKD condition aligns with and reinforces the hierarchical clustering results (Figure 6A), highlighting the activation of a specialized transcriptional network uniquely responsive to loss of Slc25a39 under glutamine-depleted conditions.

To identify a gene-function network sensitive to Slc25a39 loss under glutamine-depleted conditions (Cluster 1-2), we performed Transcriptional Regulatory Relationships Unraveled by Sentence-based Text mining (TRRUST) transcription factor target analysis^51^ for genes within Cluster 1-2, which identified Nrf2 (Nfe2l2) and Atf4 targets as the two top transcription regulators significantly affected by the combination of Slc2539 loss and glutamine withdrawal (Figure 6B). TRRUST analysis identified highly upregulated Nrf2 target genes such as *Slc7a11*, *Gclm*, *Txnip*, and *Hmox-*1 in the −QKD condition (Figure 6B). A second approach, ChIP-X-Enrichment Analysis (ChEA), also identified Nrf2 (NFE2L2) as the top enriched transcription factor (Figure S4D, Table S3). Since Nrf2 is a key regulator of cysteine and methionine metabolism, we performed KEGG pathway enrichment analysis on clusters 1-2, which revealed significant upregulation of pathways associated with selenocompound metabolism, methionine and cysteine metabolism, and one-carbon metabolism (Figure 6C). Thus, genes encoding enzymes critical for methylation cycles and detoxification, including *MAT1A*, *MTR*, and *CBS*, were upregulated. These findings indicate a broader metabolic reprogramming that reactivates one-carbon and sulfur amino acid metabolism under conditions of Slc25a39 silencing in the context of glutamine depletion (Figure 6C). We independently validated the RNA-seq results by qRT-PCR analysis of the key Nrf2 target, *Slc7a11*, and show that compared to glutamine withdrawal alone, the combination of Slc25a39 silencing and glutamine withdrawal resulted in a marked increase in *Slc7a11* mRNA expression without affecting *Nrf2* mRNA levels (Figure 6D). These results suggest that Slc25a39 might exert feedback control over Nrf2 activity under a glutamine-depleted state to establish a redox balance.

Under homeostatic conditions, Nrf2 is sequestered in cytoplasm and is continually degraded through its interaction with Keap1. In response to oxidative stress, such as that induced by complex III inhibition with Antimycin A, Nrf2 dissociates from Keap1 and translocates to the nucleus, marking the activation of the mitochondrial stress response network^42,50,52^(Figure 6E-F). However, in cells overexpressing human SLC25A39, we observed resistance to Antimycin A-induced nuclear translocation of Nrf2 despite antimycin A treatment (Figure 6E-F). Furthermore, overexpression of SLC25A39 blunted induction in *Slc7a11* expression in C2C12 myoblasts despite glutamine depletion (Figure S4E). Thus, SLC25A39 overexpression disrupts the mitochondria-to-nucleus signaling pathway, potentially by modifying the mitochondrial redox status or buffering capacity, thereby inhibiting Nrf2-dependent transcriptional activation. These findings reveal a previously unrecognized role for Slc25a39 in coordinating cellular stress responses by integrating mitochondrial metabolic signals with the transcriptional program of redox regulation, thereby communicating mitochondrial health to the nuclear machinery.

To identify cellular pathways affected by hyper-activated Nrf2 pathway in −QKD conditions, we performed gene ontology (GO) analysis, which revealed distinct gene expression patterns related to metabolic and cell cycle adaptations under glutamine-depleted (−Q) conditions compared to when glutamine depletion was combined with Slc25a39 knockdown (−QKD). Thus, whereas glycolysis was repressed under glutamine-depleted conditions (−Q), combining glutamine depletion with Slc25a39 silencing (−QKD) exhibited a notable recovery of the glycolytic transcriptional program. In addition, Slc25a39 silencing under glutamine-depleted conditions also restored the transcription program for the cell cycle, as mRNA levels of genes involved in cell cycle phase transitions were upregulated in −QKD conditions (Figure 6G, Figure S4F). These results indicate that Slc25a39 is a functional intermediate that establishes a glutamine depletion-induced quiescent-like state, and it’s silencing restores the transcriptional program of glycolysis and cell-cycle progression despite glutamine-depleted conditions. This action of Slc25a39 is likely, at least in part, via integration of mitochondrial fitness signals with Nrf2 transcriptional response to modulate glycolysis, mitochondrial membrane potential, redox balance, and cell cycle progression under nutrient-depleted conditions.

### Balanced Slc25a39 function is critical for in vitro and regenerative myogenesis

As myoblasts are acutely sensitive to metabolic and redox imbalances that influence their self-renewal and differentiation decisions^37,53^, we sought to determine the transcriptional impact of *Slc25a39* silencing on myoblast proliferation and differentiation dynamics. When analyzing multiplexed RNA-seq data for downregulated genes across individual (−Q or KD) and combined treatment groups (−QKD), we noticed a significant downregulation of genes involved in myogenesis under KD conditions, regardless of glutamine presence. In all, we identified 3,193 significantly downregulated genes (pAdj<0.05) across all conditions. Among these, 594 genes were downregulated explicitly in *Slc25a39*-silenced myoblasts (91% of DEG under KD conditions), 1,238 in the glutamine-depleted condition (83% of DEG under −Q condition), and 1361 in the combined condition (70% of DEG under −QKD condition) (Table S1). Cluster analysis of downregulated genes identified two disctinct groups with unique transcriptional responses (Figure S5A, Table 4). Clusters 6-8 included genes acutely responding to glutamine depletion either alone (−Q) or combined with Slc25a39 silencing (−QKD) but remained unaffected by Slc25a39 silencing alone (KD) (Figure S5A). Gene ontology (GO) analysis of these clusters highlighted significant downregulation of pathways related to the unfolded protein response, p53 signaling, and mTORC1 signaling, suggesting broad transcriptional repression due to glutamine deprivation and metabolic stress. Clusters 9-11 were particularly notable for their distinct response to Slc25a39 silencing. The genes in these clusters trended towards downregulation in KD myoblasts and showed variable change under glutamine depletion alone (−Q) (Figure S5A, Table S4). All three clusters feature compromised cell differentiation as a major hallmark pathway affected. Specifically, Cluster 11 was very intriguing as it featured gene sets minimally affected by glutamine depletion alone (−Q), but were dramatically downregulated by Slc25a39 knock-down alone and remained downregulated in the combined treatment conditions (−QKD). The overlapping pathway analysis of significantly downregulated genes under Slc25a39 conditions (Table S1) and from Cluster 11 revealed a pronounced and specific downregulation of the myogenesis program upon Slc25a39 knockdown (pAdj = 3.6E-24), compared to modest suppression of genes involved in myogenesis under glutamine-depleted conditions (pAdj = 5.3E-8) (Figure 7A). Thus, myoblasts are acutely sensitive to Slc25a39 loss, regardless of nutrient environment. TRRUST and ENCODE analysis revealed that MyoD1 and its downstream targets were significantly impacted by the loss of Slc25a39 in myoblasts. Thus, mRNA levels for key regulators of myogenic differentiation, including MyoD1, Mef2C, and Myog, were significantly repressed in the KD group and remained suppressed under −QKD conditions (Figure 7B). In contrast, the mRNA level of Myf5, a marker of self-renewing progenitors, was markedly upregulated, particularly in the −QKD condition (Figure 7B). This transcriptional shift underscores the role of Slc25a39 in promoting differentiation, as its silencing disrupts the expression of critical differentiation drivers while favoring the maintenance of a progenitor-like state. Under nutrient-deprived conditions, the loss of Slc25a39 exacerbates this effect, potentially pushing myoblasts toward enhancing self-renewal at the expense of differentiation.

**Figure 7.**
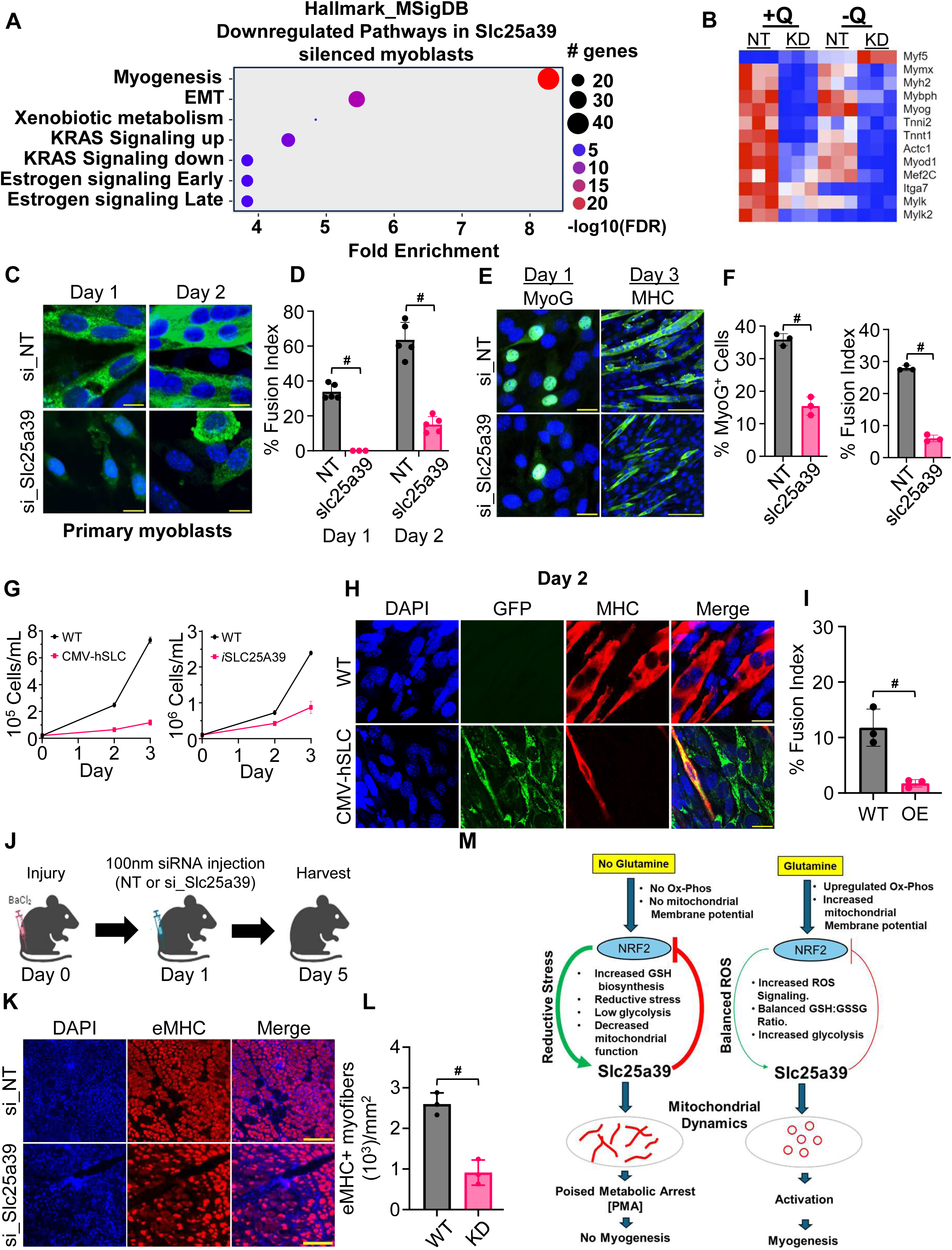
Balanced expression of Slc25a39 is necessary for proper progression through the myogenesis program *in vitro* and *in vivo*. (A) Pathway enrichment analysis of differentially expressed genes in *Slc25a39-silenced* cells, identifying affected biological pathways compared to control cells. (B) Heatmap displaying expression levels of key myogenic differentiation regulators under four conditions: control (+Q, si_NT), *Slc25a39* knockdown (+Q, si_Slc25a39), glutamine-free (−Q, si_NT), and glutamine-free *Slc25a39* knockdown (−Q, si_Slc25a39). (C) Immunofluorescence microscopy images of primary myoblasts stained for myosin heavy chain (MHC) after 24 or 48 hours of differentiation, transfected with si_NT or si_Slc25a39. Scale bar = 20 μm. (D) Quantification of fusion index (percentage of nuclei within MHC-positive multinucleated myotubes) from experiment in Figure 7C. Data represent mean ± SD from n = 5 fields per condition. Statistical significance was determined using Student’s t-test. (E) Immunofluorescence microscopy images of C2C12 cells stained for myogenin (MyoG) and MHC after 24 hours of differentiation, transfected with si_NT or si_Slc25a39. Scale bars: MyoG = 20 μm; MHC = 100 μm. (F) Quantification of fusion index and percentage of MyoG-positive cells from the experiment in Figure 7E. Data represent mean ± SD from n = 3 independent experiments. Statistical significance was determined using Student’s t-test. (G) Left: Proliferation of parental (WT) C2C12 myoblasts versus a line constitutively expressing human SLC25A39 (CMV-hSLC). Right: Proliferation of WT C2C12 versus a doxycycline-inducible tet-ON SLC25A39 line (iSLC25A39). Cells were cultured in growth medium and counted every 24 h for 72 h; WT denotes matched parental cells handled identically. Both constitutive and inducible SLC25A39 expression markedly blunts cell accumulation relative to WT. Axes reflect the observed dynamic range for each experiment. Data are mean ± SD of biological replicates. (H) Immunofluorescence images of WT and CMV-hSLC cells after 2 days of differentiation, stained for MHC (red), GFP (green), and DAPI (blue). Scale bar = 20μm. (I) Quantification of fusion index in WT and CMV-hSLC cells. Data represent mean ± SD from n = 3 independent experiments. Statistical significance was determined using Student’s t-test. (J) Schematic of in vivo experimental design: mice were subjected to muscle injury, followed by intramuscular injection of 100nM siRNA (si_NT or si_Slc25a39) the next day, and muscles were harvested on day 5 post-injury. (K) Immunohistochemistry images of injured mouse muscle sections stained for embryonic myosin heavy chain (eMHC) and DAPI, comparing si_NT and si_Slc25a39-injected muscles. Scale bar = 200 μm. (L) Quantification of muscle regeneration area, expressed as the number of eMHC-positive myofibers per mm². Data represent mean ± SD from 3 mice per group. Statistical significance was determined using Student’s t-test. (M) The proposed model illustrates how mitochondrial redox balance mediated by SLC25A39 influences myogenic differentiation and regeneration. Statistical significance is indicated as *P<0.05(*)* and *P<0.05(#)*, determined by Student’s t-test. Data are presented as mean ± S.D.

To corroborate our transcriptomic findings and further elucidate the role of Slc25a39 in myogenic differentiation in primary cells context, we silenced Slc25a39 in isolated mouse primary myoblast and monitored their differentiation performance compared to control silenced cells. Slc25a39 silencing resulted in a pronounced reduction in myosin heavy chain-positive (MHC+) cells, coupled with significantly impaired myoblast fusion, as evidenced by a marked decrease in multinucleated myotubes compared to control siRNA-treated cells (Figure 7C-D).

We further confirmed the functional role of Slc25a39 in immortalized myoblasts, as Slc25a39 knockdown in C2C12 cells attenuated Myogenin (Myog) induction, demonstrated by a reduced percentage of Myog+ cells and decreased *Myog* mRNA levels (Figure S5B), and a delayed myonuclear fusion index (Figure 7E-F). These changes were accompanied by a downregulation of key myogenic genes (Figure S5B). Together, these findings confirm that Slc25a39 is critical for myogenic progression in both primary and cultured myoblasts.

Given that Slc25a39 protein levels are tightly regulated through glutathione-dependent proteolytic degradation within mitochondria^13,14,16^ and that overexpression of SLC25A39 disrupts normal regulation of Nrf2 (Figure 6E-F, Figure S4E), we next sought to understand how its overexpression impacts myoblast function. Using clonal C2C12 cell lines engineered to overexpress human SLC25A39 under either constitutive or doxycycline-inducible conditions, we confirmed SLC25A39 expression and its mitochondrial localization via qRT-PCR and immunofluorescence (Figure S4E, Figure S5C). Overexpression of SLC25A39, regardless of the induction method, led to a significant reduction in the proliferation rate of C2C12 myoblasts (Figure 7G). Interestingly, this reduced proliferation did not correspond with an enhancement of differentiation, as SLC25A39-overexpressing myoblasts failed to progress through the early stages of the myogenic program (Figure 7H-I). These results underscore the importance of precise proteostatic regulation of Slc25a39 in coordinating myoblast function. While its silencing disrupts differentiation and fusion, overexpression impairs proliferation and blocks differentiation, suggesting that tight control of Slc25a39 expression is essential for the temporal and functional progression of myoblasts through the myogenic differentiation program.

While the in vitro myogenesis program shares core molecular machinery with in vivo regenerative myogenesis, it remains unclear whether mitochondrial dysfunction, particularly due to Slc25a39 loss, impacts the progression of regenerative myogenesis in vivo. To investigate this, we induced muscle injury in mice using BaCl₂, which triggers myonecrosis and initiates MuSCs-driven regeneration. Twenty-four hours post-injury, control or Slc25a39 siRNA was injected into the injured muscle, and regeneration was assessed after four days (Figure 7J). In control siRNA-treated muscles, we observed robust regeneration, as indicated by the emergence of embryonic MHC (eMHC+)-positive myofibers, a hallmark of regenerative myogenesis. In contrast, Slc25a39 silencing resulted in a profound defect in regenerative myogenesis, evidenced by a significant reduction in percent eMHC+ regenerating myofibers (Figure 7K-L). These findings establish Slc25a39 as a critical regulator of myoblast differentiation and regenerative myogenesis, operating within the mitochondrial glutamine-Nrf2 axis to maintain cellular redox balance and metabolic flexibility, which are essential for muscle repair.

## Discussion

Our study uncovers a redox-based mechanism to establish muscle stem cell quiescence in the absence of mitochondrial glutamine metabolism (Figure 7M). The mitochondrial GSH transporter Slc25a39 is central to this process, which functions as a redox-sensitive mediator, linking mitochondrial health to nuclear transcriptional responses via Nrf2 signaling and regulating the balance between reductive and oxidative stress. Under glutamine-depleted conditions, Nrf2 activation and increased Slc25a39 expression drive mitochondrial remodeling, glycolysis repression, loss of mitochondrial membrane potential, and decreased oxygen consumption, ultimately creating a reductive stress environment. Building on the evidence that Slc25a39 silencing reverses glycolytic repression and restores membrane potential in myoblasts under glutamine-starved conditions, we propose that sustained Slc25a39 expression maintains a reductive stress environment via feedback regulation of Nrf2. This mechanism facilitates metabolic adaptation to nutrient deprivation, promoting cell cycle withdrawal and the transition to a PMA state (Figure 7M). Conversely, mitochondrial activation through glutaminolysis establishes a balance between oxidative and reductive processes, enabling ROS signaling essential for pro-growth pathways and restoring normal regulation of Slc25a39 and Nrf2. As a mitochondrial GSH importer, these findings establish Slc25a39-mediated GSH metabolism as a critical, previously underappreciated factor in establishing the reductive environment necessary for acquiring a PMA state under glutamine-depleted conditions.

### Glutamine depletion induced cell state transition

In contrast to MRCs, which arise during the differentiation process, the glutamine limitation in our study was applied to actively proliferating, uncommitted myoblast. Yet, it drove cells into a quiescent-like state, which is molecularly characterized as PMA. While the molecular mechanisms that govern the bifurcation of differentiating myoblasts into either fusogenic or reserve cell fates remain poorly understood, our findings raise the possibility that reduced glutamine uptake or mitochondrial glutamine metabolism may underlie MRC emergence. In this context, our results provide a mechanistic framework for understanding how metabolic signals— specifically glutamine availability—can regulate the transition between various proliferative and non-proliferative states amongst myoblast progenitors.

### Glutamine Metabolism and the Coordination of Mitochondrial Dynamics and Glycolysis

Our data highlight the crucial role of glutamine metabolism in reconfiguring mitochondrial networks and stimulating glycolytic pathways during myoblast proliferation. Glutamine-depleted myoblasts retained elongated mitochondria with reduced mitochondrial membrane potential, oxygen consumption, and ROS production, indicating a state of quiescence reminiscent of resting muscle stem cells (Figure 1A–G, Figure S1A-C). This suggests that glutamine metabolism provides essential signals for mitochondrial fission and remodeling, a prerequisite for the activation and differentiation of energy-demanding proliferating myoblasts. This metabolic adaptation is necessary for promoting mitochondrial function and maintaining glycolytic gene expression, as evidenced by the repression of key glycolytic genes such as *Pgk1*, *Pgm2*, and *Me2* upon glutamine depletion (Figure 1K–L). These findings support a model in which glutamine serves as a central metabolic regulator in myoblasts, maintaining mitochondrial dynamics and metabolic flux through glycolysis to support the transition from quiescence to an activated myogenic state.

The partial rescue of mitochondrial fragmentation and myogenic function by α-ketoglutarate (α-KG) supplementation (Figure 2B–C) underscores the importance of glutamine-derived metabolites in regulating mitochondrial structure and energy metabolism. The downstream metabolism of α-KG, which is central to redox buffering and GSH regeneration, further points to the integration of mitochondrial and cytosolic metabolic networks in adapting to environmental changes. Inhibition studies targeting GOT1 (Figure 2D) and Glud1 reinforce this concept, revealing that glutamine-derived α-KG provides essential bioenergetic substrates and maintains mitochondrial GSH levels, which are critical for sustaining cellular redox homeostasis during MuSCs activation. These findings indicate that glutamine metabolism facilitates mitochondrial remodeling and transcriptionally regulates metabolic pathways to ensure bioenergetic and biosynthetic demands are met.

### Slc25a39 as an Nrf2-Dependent Effector In Glutamine-Deprived Myoblasts

Our transcriptomic analyses reveal Slc25a39 as a downstream target of Nrf2, which is activated in response to glutamine depletion and oxidative stress and functions as a key regulator of antioxidant defense genes, including those involved in GSH biosynthesis and transport (Figure 3F, Figure 4C). The induction of Slc25a39 in glutamine-depleted myoblasts was shown to be Nrf2-dependent, as Nrf2 knockdown abolished Slc25a39 upregulation under both glutamine-depleted and oxidative stress conditions (Figure 4D). This finding expands the known repertoire of Nrf2 targets and highlights Slc25a39 as a critical effector within the Nrf2-driven antioxidant response. The glutamine-depleted environment, which suppresses glycolysis and elevates reductive stress, thus engages Slc25a39 to import GSH into mitochondria, mitigating the potential oxidative damage that may arise under nutrient-limiting conditions.

The upregulation of Slc25a39 and other Nrf2 target genes, such as Slc7a11 and Hmox1, in response to glutamine depletion (Figure 4B–C) suggests a broader adaptive response to low nutrient availability aimed at fortifying mitochondrial redox defenses. This Nrf2-driven transcriptional program is crucial for maintaining mitochondrial integrity and supporting myogenesis, as oxidative stress-induced damage would otherwise compromise the cellular machinery required for proliferation and differentiation. Our findings also imply that Nrf2, through Slc25a39, supports mitochondrial redox buffering, which, in turn, facilitates proper muscle stem cell function under fluctuating nutrient conditions.

### Redox Sensing by Slc25a39 Enables Feedback Control on Nrf2 Activity

Our findings reveal a complex, bidirectional interaction between Slc25a39 and Nrf2, where Slc25a39 not only acts as a downstream target but also regulates Nrf2 activity by controlling mitochondrial GSH levels. In glutamine-depleted cells, silencing Slc25a39 led to heightened expression of Nrf2 target genes, including Slc7a11, indicating a feedback loop where mitochondrial GSH status communicates with nuclear antioxidant responses (Figure 6B and 6D). This feedback mechanism suggests that Slc25a39, as a mitochondrial redox sensor, modulates Nrf2 activity, thereby enabling cells to adjust to metabolic stress and oxidative fluctuations dynamically.

Overexpression of Slc25a39 disrupted Nrf2 activation in response to oxidative stress, as evidenced by reduced nuclear translocation of Nrf2 and blunted expression of downstream targets, even under antimycin A treatment (Figure 6E-F). This suggests that Slc25a39, by buffering mitochondrial redox conditions, limits Nrf2’s nuclear signaling under oxidative stress, highlighting a critical function of Slc25a39 in coordinating mitochondrial and nuclear antioxidant defenses. These findings underscore the dual role of Slc25a39 in both sensing mitochondrial redox status and relaying this information to the Nrf2 pathway, positioning it as a central mediator in balancing oxidative and reductive stress.

### Requirement of Mitochondrial GSH Import for Myogenesis

The mitochondrial import of GSH via Slc25a39 emerges as an essential factor in myogenic progression, as both loss and gain of Slc25a39 function disrupted myogenesis in vitro and vivo. Silencing Slc25a39 impaired mitochondrial membrane potential and function and inhibited the expression of myogenic genes, including MyoD1, Myog, and Mef2C (Figure 5E–F, Figure 7B). The observed mitochondrial fragmentation in glutamine-depleted cells upon Slc25a39 silencing (Figure 5A-B) further underscores the importance of mitochondrial GSH in maintaining structural integrity and bioenergetic function during myogenesis.

Interestingly, overexpression of human SLC25A39 also resulted in inhibition of myoblast proliferation and inhibited myogenesis of C2C12 myoblasts (Figure 7G–I). These results indicate the need for precise regulation of Slc25a39 activity, as insufficient and excessive GSH import could disturb mitochondrial function and cellular differentiation processes. This tightly regulated import system ensures that mitochondrial GSH levels are optimized for redox buffering without overburdening the system, highlighting the critical role of Slc25a39 in maintaining a balanced redox environment conducive to myogenic differentiation.

### Role of Mitochondrial GSH in Regulating Glycolytic Gene Expression

Our findings suggest that mitochondrial GSH is regulatory in glycolytic gene expression, serving as a feedback mechanism to mitigate oxidative damage. Under glutamine-depleted conditions, elevated mitochondrial GSH establishes a reductive stress environment that inhibits glycolytic transcription (Figure 6G). This suppression of glycolysis is relieved by Slc25a39 knockdown, restoring glycolytic gene expression and mitochondrial membrane potential, which may reflect an adaptive response to restore cellular redox balance and energy production. Thus, mitochondrial GSH appears to act as a modulator of glycolysis under redox-stress conditions, ensuring that cellular metabolism aligns with mitochondrial redox requirements.

The modulation of Nrf2 activity and glycolytic gene expression by mitochondrial GSH highlights a protective mechanism whereby myoblasts prevent oxidative damage, balancing energy production with antioxidant capacity. This control of glycolysis through mitochondrial GSH import may represent a novel strategy for cells to safeguard redox balance, allowing efficient myoblast proliferation and differentiation in response to nutrient stress.

## Acknowledgement

We thank members of the Kikani lab for the discussion and critical reading of the manuscript. This work was supported by funding from NIAMS to C.K.(R01AR073906) and from NIGMS to J.S (R35GM130349).

## Experimental Model and Subject Details

### Animals

C57BL/6 mice (8–12 weeks old) were used for muscle regeneration experiments and for isolating muscle stem cells (MuSCs). All mice were housed under standard conditions with a 12-hour light/dark cycle, temperature of 22 ± 2°C, and ad libitum access to food and water. All animal procedures were approved by the Institutional Animal Care and Use Committee (IACUC) of the University of Kentucky (protocol number 2022-3317).

### Muscle Stem Cell Isolation and Culture

Muscle stem cells (MuSCs) were isolated using a two-step magnetic-activated cell sorting (MACS) protocol, initially depleting non-MuSCs by negative selection, followed by specific purification of integrin α7-positive (Integrin α7⁺) MuSCs. Briefly, tibialis anterior (TA) muscles were harvested from 8 weeks old C57BL/6 mice, minced, and enzymatically digested in a solution containing 0.2% Collagenase Type II and 0.1% Dispase II at 37°C for 60–90 minutes with intermittent mixing. The digested tissue was triturated to dissociate cells, and enzymes were removed by washing with cold serum-free DMEM+F12. The cell suspension was filtered through a 70 µm cell strainer and centrifuged at 300 × g for 5 minutes. The cell pellet was washed, resuspended in MACS buffer (PBS containing 0.5% bovine serum albumin [BSA] and 2 mM EDTA), and processed using a satellite cell isolation kit (Miltenyi, Catalog 130-104-268) to remove non-MuSCs. The flow-through containing MuSCs was then incubated with magnetic beads conjugated to an antibody against integrin α7 (Miltenyi, Catalog 130-104-261), and passed through a MACS column according to the manufacturer’s instructions. Labeled MuSCs were retained in the magnetic field, while unbound cells were washed away. The enriched integrin α7⁺ MuSCs were eluted, washed, and cultured in myoblast proliferation media (see below).

### Cell Culture

Isolated MuSCs were maintained (now referred to as primary mouse myoblasts) in proliferation medium consisting of DMEM/F12 supplemented with 20% fetal bovine serum (FBS), 10% horse serum, 1.5 ng/mL basic fibroblast growth factor (bFGF), and 1% penicillin-streptomycin (Pen/Strep). C2C12 myoblasts and HEK293T cells were cultured in high-glucose DMEM (4.5 g/L glucose) supplemented with 10% FBS, 2 mM L-glutamine, 1 mM sodium pyruvate, and 1% Pen/Strep at approximately 60% confluence. Cells were maintained at 37°C in a humidified atmosphere with 5% CO₂.

For experiments involving glutamine depletion, proliferation media was exchanged with glutamine-free high-glucose DMEM containing sodium pyruvate. Control cells were supplemented with 2 mM L-glutamine to match standard culture conditions. Differentiation medium for primary myoblasts and C2C12 cells consisted of DMEM supplemented with 2% horse serum, 1 mM sodium pyruvate, and 1% Pen/Strep. For siRNA knockdown experiments, siRNA duplexes complexed with Lipofectamine RNAiMAX reagent (Thermo Fisher Scientific) in Opti-MEM reduced-serum medium were added to cells in maintenance medium.

### Cloning of Human SLC25A39 and MitoTag-mCherry

Human SLC25A39 was amplified from cDNA derived from HEK293T cells and cloned into either the pQCXIP-GFP vector or a custom pQCXIP-Tet-ON-Flag-HA-T2A-dsRED-Puro-T2A-rTTA vector generated in our laboratory. Whole plasmids were sequenced using nanopore sequencing technology to verify insert orientation and lack of inadvertent mutations. An mCherry-tagged mitochondrial marker (MitoTag-mCherry) was generated by PCR-based subcloning of the 3X-HA-OMP25 sequence from the 3X-HA-OMP25-EGFP construct (pMXs-3XHA-EGFP-OMP25 was a gift from David Sabatini; Addgene plasmid #83356).

### Metabolic Pathway Inhibitor Treatments

To analyze pathways downstream of glutamine metabolism, various metabolic inhibitors were introduced. After 24 hours of proliferation in a glutamine-containing medium, cells were exposed to either glutamine-depleted DMEM (−Q) or glutamine-depleted DMEM supplemented with 2 mM L-glutamine (+Q). Modulators of metabolic pathways, including 10 µM BPTES, 10 µM UK5099, 3 mM dimethyl α-ketoglutarate (αKG), 5 mM sodium succinate (SUCC), 10 µM R162, 4 µM GOT1 inhibitor (GOT1i), 1 mM buthionine sulfoximine (BSO), 500 µM reduced glutathione ethyl ester (GSH-EE), 2 µM mitoCDNB (mCDNB), or 100 µM diamide, were added concurrently with glutamine depletion for an additional 24 hours.

To assess C2C12 cell viability under glutamine-depleted conditions, cells were cultured in 10 cm plates in normal growth medium for 24 hours, followed by a change to either normal growth medium or glutamine-free medium containing high or low serum, with either 1 mM BSO or vehicle (water) for 24 hours. Cell viability was quantified using the Trypan Blue exclusion assay.

Cell proliferation rates were measured daily using the Invitrogen Countess II FL Automated Cell Counter. GFP-vector control C2C12 cells, CMV-hSLC25A39-GFP overexpressing cells, and tet-on inducible SLC25A39-overexpressing (iSLC25A39) cells were seeded in 10 cm plates at a density of 1 × 10⁵ cells per plate. For induction of SLC25A39 expression, doxycycline was added at a final concentration of 2 µg/mL at the time of seeding. Cells were grown and counted daily.

### siRNA Knockdown

ON-TARGETplus Non-targeting Control siRNA or target-specific ON-TARGETplus SMARTpool siRNA (50 nM; Dharmacon) was diluted in Opti-MEM reduced-serum medium (Thermo Fisher Scientific) and complexed with Lipofectamine RNAiMAX reagent according to the manufacturer’s instructions. The siRNA–Lipofectamine complexes were added to cells in suspension immediately after trypsinization. Cells were allowed to take up siRNA and were cultured for 24 hours prior to subsequent treatments.

### Myoblast differentiation

Differentiation was induced when cells reached approximately 90% confluence by replacing the growth medium with differentiation medium (DMEM containing 2% horse serum) for the indicated number of days. At the end of the differentiation time course, cells were fixed and stained for specific antibodies as described below. The fusion index was calculated as the percentage of nuclei within multinucleated myotubes over the total number of nuclei within the field of view.

## Method Details

### Cell Line Generation

Stable C2C12 cell lines expressing CMV-SLC25A39-GFP, inducible SLC25A39 (iSLC25A39), or MitoTag-mCherry were generated using retroviral transduction. Transduced cells were selected using puromycin (2 µg/mL) until non-transduced control cells were eliminated.

### Muscle injury and in vivo siRNA delivery

All procedures adhered to approved institutional protocols and animal-welfare guidelines. WT C57BL6 mice (n=3 per each control and Slc25a39 knock-down conditions) were anesthetized with continuous-flow isoflurane (2–3% induction, 1–2% maintenance in oxygen). Under aseptic conditions, the skin over the tibialis anterior (TA) was sterilized, and 50 µL of sterile 1.2% BaCl₂ (or PBS control) was injected into the mid-belly of the TA using a 29-g insulin syringe, advancing the needle along the fiber axis to minimize leakage. This dose reliably activates MuSCs without excessive necrosis^21^. Animals were monitored until recovery. For in vivo knockdown at 1 day post-injury (1 DPI), chemically synthesized siRNA duplexes targeting Slc25a39 (or non-targeting control) were prepared fresh at 1 DPI at a final concentration of 100 nM in sterile PBS and complexed with RNAiMAX (Thermo Fisher) for 10–15 min immediately before injection. Fifty microliters of the siRNA–RNAiMAX complex was then injected into the mid-belly of each TA (bilateral) using a 29-g syringe. Mice received routine post-procedure monitoring. qRT-PCR at 5 DPI verified knockdown (∼60% reduction in Slc25a39 mRNA).

At 5 days post-injury (5 DPI), TAs were harvested to capture the early regenerative response. Muscles were embedded in OCT, frozen in isopentane pre-chilled on liquid nitrogen, and cryosectioned (8–10 µm). Sections were processed for embryonic myosin heavy chain (eMHC) immunostaining (with DAPI) to quantify early regeneration. Quantification was performed blinded to condition.

### Immunofluorescence

Cells were grown on glass coverslips in 24-well plates. Cells were fixed with 4% ice-cold paraformaldehyde in PBS containing Ca²⁺ and Mg²⁺ at room temperature for 20 minutes. Following fixation, cells were washed three times with PBS and permeabilized with 0.2% Triton X-100 in PBS for 10 minutes. Cells were then blocked with 10% normal goat serum at room temperature for 1 hour. Primary antibodies diluted in blocking buffer were added to the cells and incubated overnight at 4°C in a humidified chamber. The following day, cells were washed three times with PBS and incubated with fluorescently labeled secondary antibodies for 1 hour at room temperature in the dark. After washing, nuclei were stained with DAPI for 15 minutes at room temperature in the dark. Coverslips were mounted onto slides using an antifade mounting medium and allowed to cure overnight in the dark.

### Mitochondrial Length Measurement

To measure mitochondrial length, freshly isolated primary myoblasts were cultured in either normal growth medium or glutamine-free medium for 24 hours. Mitochondrial length was assessed using MitoView Green (Biotium), a mitochondrial-specific dye, in live-cell imaging. For C2C12 cells, stable MitoTag-mCherry–expressing cells were seeded onto coverslips and cultured similarly. After treatment, cells were fixed and processed for immunofluorescence as described above. In experiments involving chemical treatments, compounds were added concurrently with glutamine withdrawal or as indicated. Mitochondrial length was measured using FIJI software (ImageJ). Ten individual cells per condition were imaged, and the lengths of all mitochondria within each cell were measured, averaged, and plotted using GraphPad Prism.

### Quantitative Real-Time PCR (qRT-PCR) Analysis

Total RNA was extracted from cells using the RNeasy Kit (Qiagen) according to the manufacturer’s instructions. RNA concentration and purity were assessed using an Implen NP80 NanoPhotometer. cDNA was synthesized from 2 µg of total RNA using the High-Capacity cDNA Reverse Transcription Kit (Thermo Fisher Scientific). qRT-PCR was performed on a QuantStudio 6 Pro system using SYBR Green Master Mix (QuantaBio). Primer sequences are listed in Supplemental Table S3. All experiments were performed in biological triplicate. Gene expression levels were normalized to the housekeeping gene 18S rRNA. Statistical analysis was performed using a paired Student’s t-test, with *p*<0.05 considered statistically significant. Error bars represent the mean ± standard deviation (SD).

For RNA sequencing, C2C12 cells were cultured in glutamine-replete or glutamine-depleted proliferation medium under control or mouse Slc25a39-silenced conditions. Total RNA was extracted as described above, and sequencing libraries were prepared using the TruSeq PolyA Enrichment Library Prep Kit (Illumina) and sequenced at Azenta Inc.

### Glutathione (GSH) Measurement Assay

Total glutathione levels (GSH + GSSG) were measured using the GSH/GSSG Assay Kit (Abcam) according to the manufacturer’s instructions. C2C12 cells were cultured in 6 cm plates under control and glutamine-depleted conditions as described above. Samples were prepared following the kit protocol, and GSH levels were quantified using a BioTek Synergy MX Microplate Reader. Data were analyzed using BioTek Gen5 Software.

### Mitochondrial Functional Tests

Oxygen consumption rate (OCR) and extracellular acidification rate (ECAR) were measured using an Agilent Seahorse XFe96 Analyzer. Cells were seeded into poly-D-lysine–coated 96-well Seahorse cell culture plates and cultured under the specified conditions. Assays were performed in collaboration with the Redox Metabolism Shared Resource Facility. During compound injections, glucose-free and glutamine-free media were used as appropriate for each condition.

Mitochondrial membrane potential (ΔΨm) was measured using the JC-1 dye (Thermo Fisher Scientific) in live cells. Cells were cultured under control or glutamine-depleted conditions for 24 hours. JC-1 dye was added to each well at a final concentration of 2 µM and incubated for 15 minutes at 37°C in the dark. Fluorescence was measured using excitation/emission wavelengths of 488 nm/530 nm (monomers, green) and 568 nm/590 nm (J-aggregates, red). The ratio of red to green fluorescence intensity was calculated using FIJI software and plotted using GraphPad Prism.

### RNA-seq Analysis

Raw RNA-seq data were obtained as FASTQ files and subjected to quality control using FastQC to assess sequencing quality, adapter contamination, and sequence duplication levels. Low-quality bases and adapter sequences were trimmed using Trimmomatic. Cleaned reads were aligned to the *Mus musculus* reference genome (GRCm39) using HISAT2 v2.2.1. Gene expression levels were quantified using StringTie v2.2.0, and differential gene expression analysis was performed using DESeq2 in R. Genes with an adjusted *p*-value (false discovery rate, FDR) of less than 0.05 were considered significantly differentially expressed. Pathway enrichment analyses were conducted using Gene Set Enrichment Analysis (GSEA) and Enrichr to identify biological processes and signaling pathways affected by the experimental conditions.

