## Supplemental Figures for "Glutamine-Dependent Slc25a39–Nrf2 Axis Couples Mitochondrial Dynamics with Metabolic Reprogramming to Establish Myogenic Commitment"

### Supplementary Material for

### Coordination of mitochondrial dynamics with transcriptional program of myogenesis through glutamine-anchored reciprocal Slc25a39–Nrf2 retrograde signaling axis

**Kelly B et al.,**

**Figure S1-S5**

**Supplementary Figure Legends**

**Figure S1. Glutamine availability regulates myoblast proliferation and metabolic programs.**
(A) Proliferation rate of primary myoblasts cultured in the presence (+Q) or absence (–Q) of glutamine. The arrow marks the switch to glutamine-free conditions.
(B) Proliferating primary myoblasts cultured with or without glutamine were pulsed with 10 μM BrdU for 24 h and analyzed by immunofluorescence using anti-BrdU and anti-Pax7 antibodies.
(C) Quantification of $Pax7^+/BrdU^+$ cells from three independent experiments in (B). Data represent mean ± S.D. *P < 0.05.
(D-E) PMA cells can reenter the cell cycle upon glutamine addition: C2C12 myoblasts stably expressing MitoTag-mCherry were glutamine-starved for 72 hours to ensure establishment of the PMA state, after which, 2mM Glutamine was added along with 10μM BrdU for 48 hours. Cells were fixed and stained using an anti-BrdU antibody. Scale bar = 40μM. Quantification in (E) represents averages from three independent experiments. Error bars represent ± S.D.
(D) Gene Set Enrichment Analysis (GSEA) from RNA-seq data of C2C12 cells cultured in +Q or –Q media.
(E) Heatmap of central carbon metabolites from metabolomic profiling of C2C12 cells in +Q or –Q media.
(F) Extracellular acidification rate (ECAR) of C2C12 cells cultured in glucose-replete (+Gluc) or glucose-free (–Gluc) media with BPTES or vehicle (DMSO) during ATP-linked respiration ($n$ = 18).
(G) ECAR of C2C12 cells cultured in +Q or –Q media during basal respiration ($n$ = 6). *P < 0.05; #P < 0.01. Data are shown as mean ± S.D.

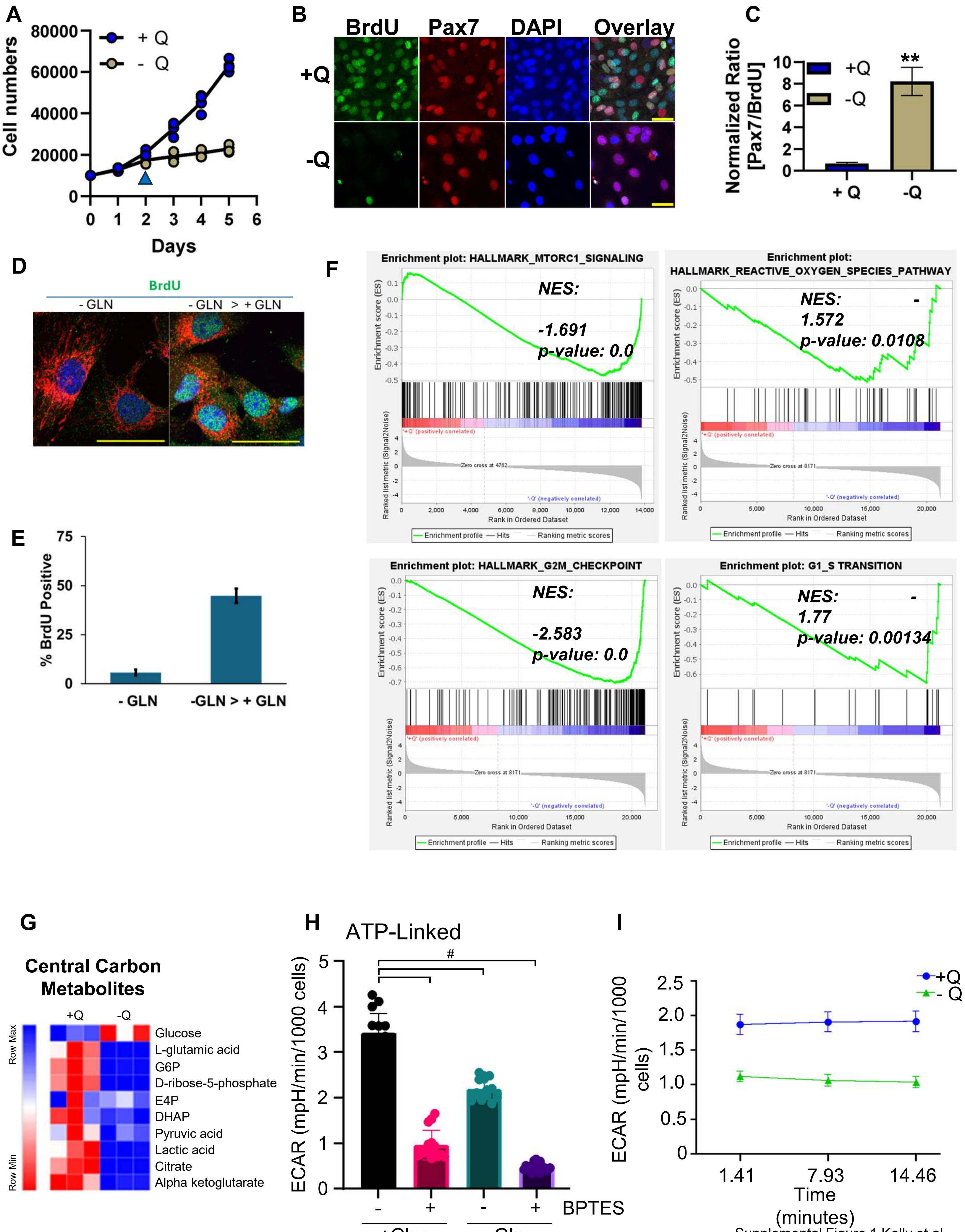


Supplemental Figure 1 Kelly et al.

**Figure S2. Identification of metabolic intermediates in glutamine metabolism regulating mitochondrial length in myoblasts.**

(A) Immunofluorescence microscopy of C2C12 cells cultured in differentiation media for 2 days in +Q or –Q conditions, with α-ketoglutarate (αKG), succinate, or vehicle (Veh), stained for myosin heavy chain (MHC) and DAPI. Scale bar = 40 μm.

(B) Quantification of fusion index ($n$ = 5).

(C) Immunofluorescence microscopy of C2C12 cells cultured in differentiation media for 2 days in +Q or –Q conditions, with αKG, GOT1 inhibitor (GOT1i), R162, or vehicle (Veh), stained for MHC and DAPI. Scale bar = 20 μm.

(D) Quantification of multinucleated myotubes relative to mononucleated myotubes ($n$ = 5).

(E) Glutathione (GSH) assay of proliferating C2C12 cells cultured in +Q or –Q media ($n$ = 3). *P < 0.05; #P < 0.01. Data are shown as mean ± S.D.

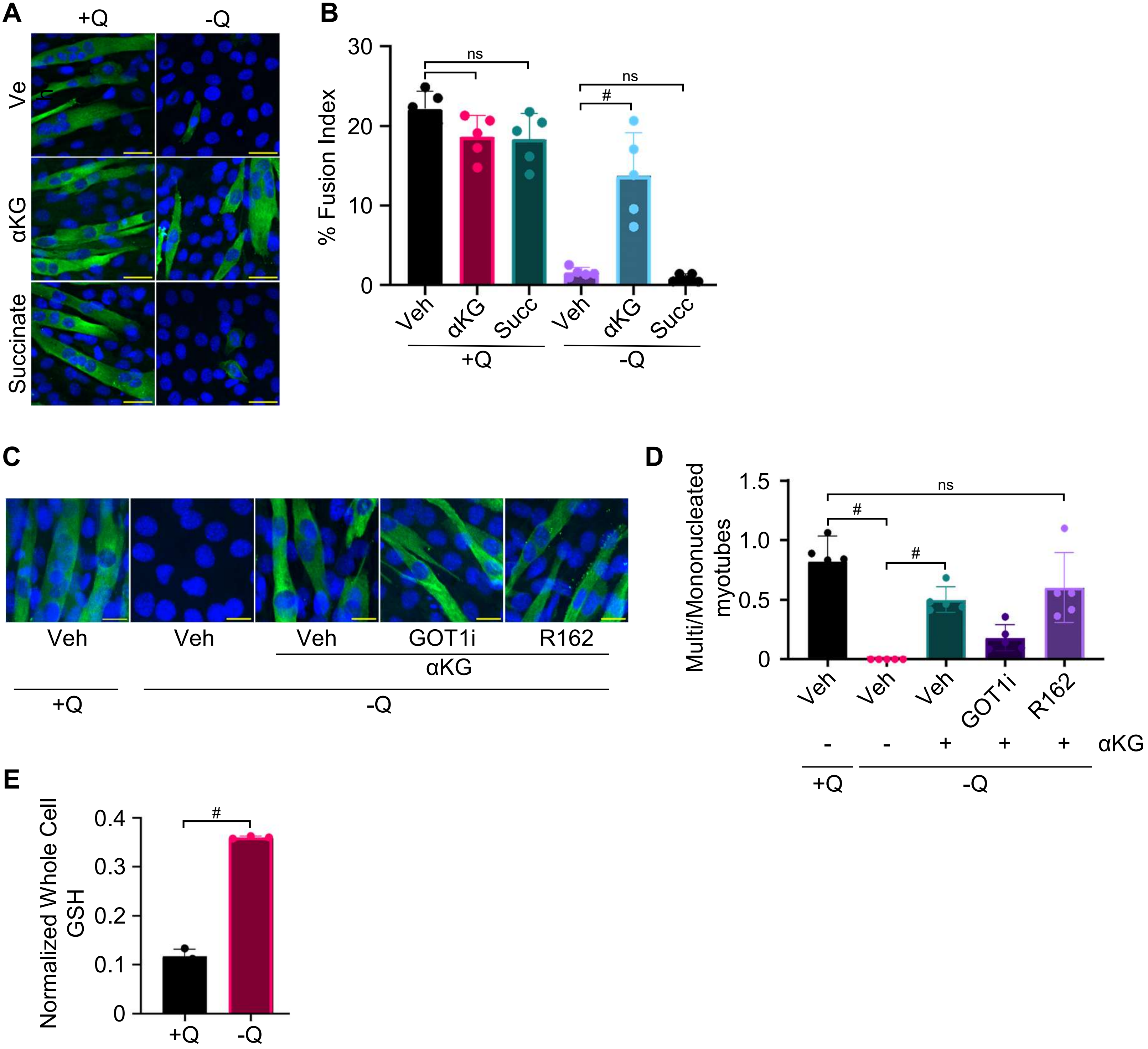


Supplemental Figure 2 Kelly et al.

**Figure S3. Transcriptional induction of Atf3 and Atf4 pathways under glutamine-depleted conditions.**
(A) Fragments per kilobase of transcript per million mapped reads (FPKM) values of *Chac1*, *Atf3*, and *Atf4* from RNA-seq analysis.

A

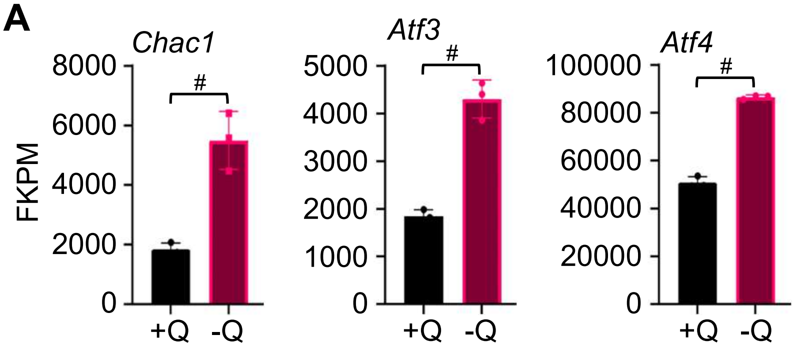
Chac1
Atf3
Atf4
FKPM
8000
6000
4000
2000
0
5000
4000
3000
2000
1000
0
100000
80000
60000
40000
20000
0
#
#
#
+Q
-Q
+Q
-Q
+Q
-Q

**Figure S4. Combinatorial RNA-seq analyses identify feedback regulation of Nrf2 by Slc25a39.**

(A) Principal component analysis comparing control (+Q), glutamine-free (–Q), *Slc25a39* siRNA knockdown (KD), and *Slc25a39* KD in glutamine-free media (–QKD).
(B) k-means clustering of transcriptomic profiles across the same conditions.
(C) Heatmap of NRF2/ATF4 pathway genes from RNA-seq data of control (+Q), glutamine-free control (–Q), *Slc25a39* KD (+Q), and *Slc25a39* KD (–Q).
(D) ChIP-X Enrichment Analysis (ChEA) of *Slc25a39* KD C2C12 cells in –Q conditions.
(E) qRT-PCR of proliferating WT C2C12 and CMV-hSLC cells ($n$ = 3).
(F) Gene Set Enrichment Analysis of the hallmark glycolysis gene set comparing –Q vs +Q control cells and –QKD vs –Q control cells. *$P < 0.05$; #$P < 0.01$. Data are shown as mean ± S.D.

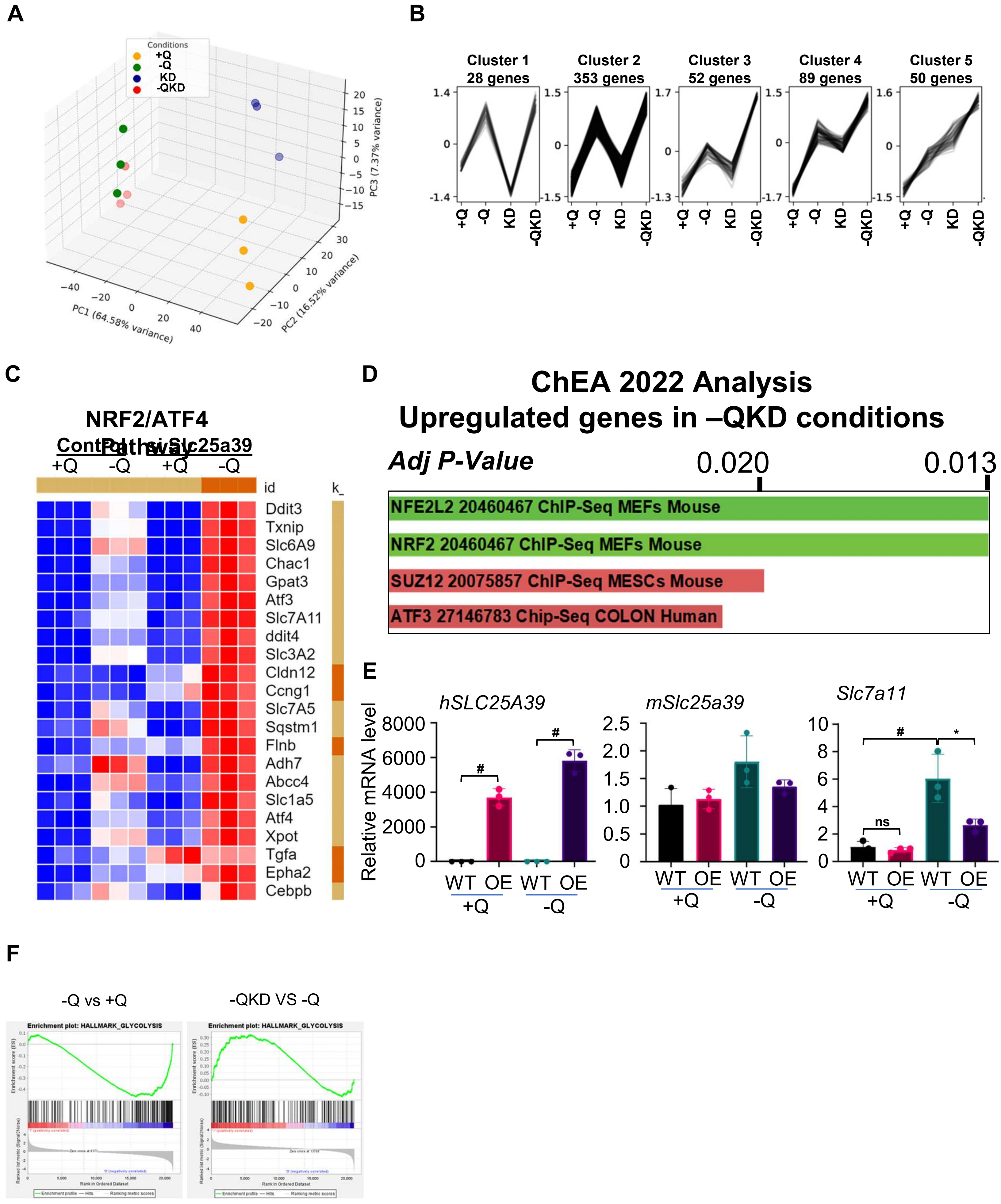


Supplemental Figure 4 Kelly et al.

**Figure S5. Regulation of myogenesis by Slc25a39.**
(A) k-means clustering analysis of cells cultured in control (+Q), glutamine-free (–Q), *Slc25a39* KD (+Q), and *Slc25a39* KD (–Q) conditions.
(B) qRT-PCR of C2C12 cells cultured in differentiation media for 0 or 2 days with non-targeting siRNA (Con) or *Slc25a39* siRNA (KD) ($n$ = 3).
(C) Immunofluorescence microscopy of CMV-hSLC cells showing GFP and DAPI. Scale bar = 20 μm. *$P < 0.05$; #$P < 0.01$. Data are shown as mean ± S.D.

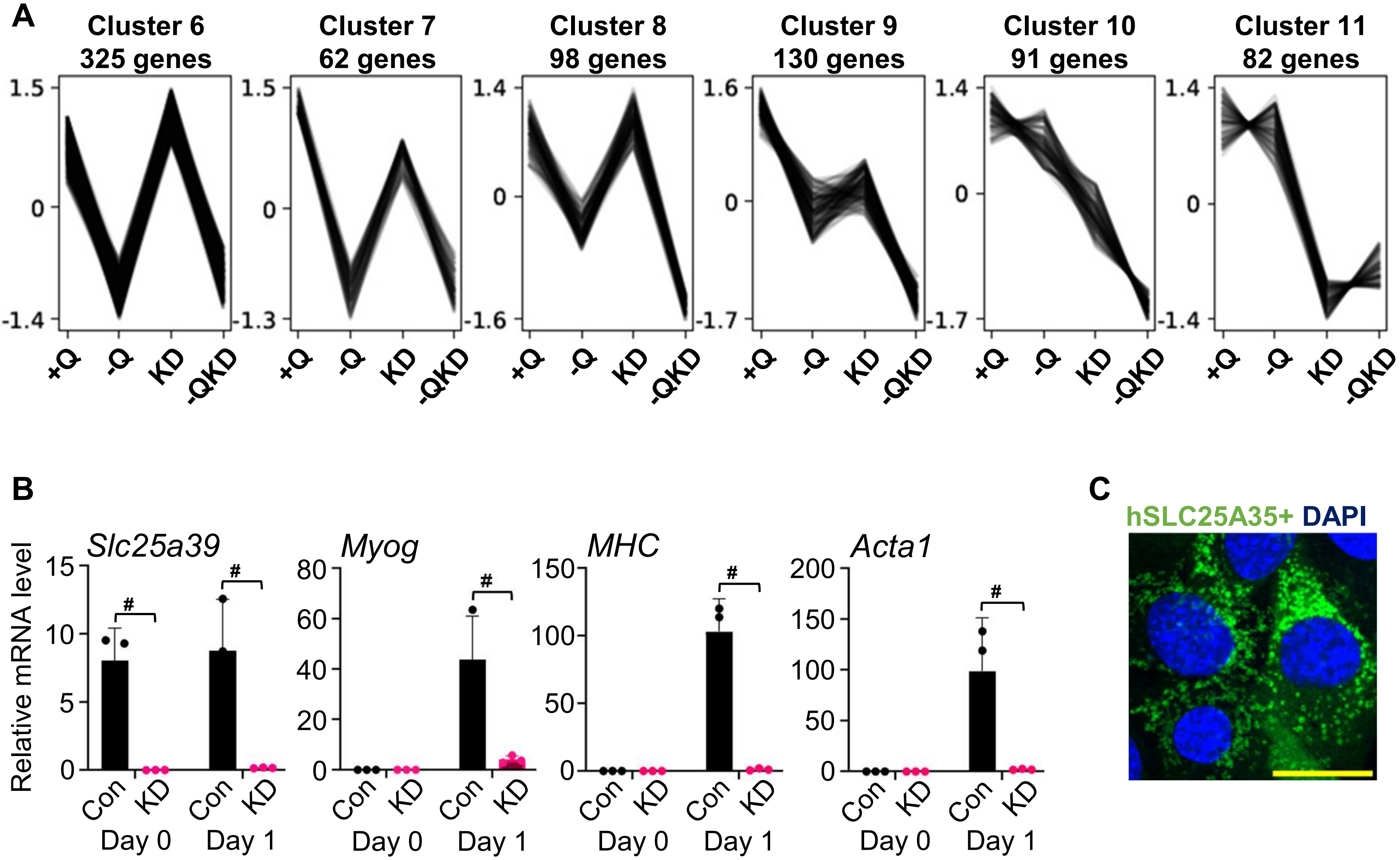
A
Cluster 6
325 genes
Cluster 7
62 genes
Cluster 8
98 genes
Cluster 9
130 genes
Cluster 10
91 genes
Cluster 11
82 genes
+Q
-Q
KD
-QKD
B
Slc25a39
Myog
MHC
Acta1
Relative mRNA level
Con
KD
Day 0
Day 1
C
hSLC25A35+ DAPI
